# A translation-independent directed evolution strategy to engineer aminoacyl-tRNA synthetases

**DOI:** 10.1101/2023.12.13.571473

**Authors:** Chintan Soni, Noam Prywes, Matthew Hall, David F. Savage, Alanna Schepartz, Abhishek Chatterjee

## Abstract

Using directed evolution, engineered aminoacyl-tRNA synthetases (aaRS) have been developed that enable co-translational incorporation of numerous noncanonical amino acids (ncAAs) into proteins in living cells. Until now, the selection of such novel aaRS mutants has relied on coupling their activity to the expression of a reporter protein with a selectable phenotype. However, such translation-dependent selection schemes are incompatible with exotic monomers that diverge structurally from canonical α-amino acids and are suboptimal substrates for the ribosome. To enable the ribosomal incorporation of such exotic monomers, a two-step solution is needed: A) Engineering an aaRS to acylate its cognate tRNA with the exotic monomer, without relying on ribosomal translation as a readout, and B) Subsequent engineering of the ribosome to accept the resulting acylated tRNA for translation. Here, we report a platform for aaRS engineering that directly selects for tRNA-acylation without ribosomal translation (START). In START, each distinct aaRS mutant is correlated to a cognate tRNA containing a unique sequence barcode. Acylation by an active aaRS mutant protects the associated barcode-containing tRNAs from an oxidative treatment designed to damage the 3′-terminus of the uncharged tRNAs. Sequencing of these surviving barcode-containing tRNAs is then used to reveal the identity of aaRS mutants that acylated the correlated tRNA sequences. The efficacy of START was demonstrated by identifying novel mutants of the *M. alvus* pyrrolysyl-tRNA synthetase from a naïve library that charge noncanonical amino acids.

## Introduction

Over the last two decades, the genetic code of cells in various domains of life has been expanded to include hundreds of noncanonical amino acids (ncAAs) with diverse chemical structures.^1–4^ This expansion has been accomplished by developing engineered aminoacyl-tRNA synthetase/tRNA pairs that selectively incorporate various ncAAs in response to a unique codon (e.g., a repurposed nonsense codon).^1–5^ The ability to site-specifically incorporate enabling ncAAs into proteins using this technology has unlocked powerful new ways to probe and manipulate protein function for both basic science and biotechnology applications.^1–3, 6, 7^

Despite such exciting progress, this technology has been largely restricted to the incorporation of simple L-α-amino acids and, to a lesser extent, some α-hydroxy acids.^8–12^ Pioneering early work using *in vitro* translation systems has demonstrated that the ribosome has the potential to accept a much wider variety of monomers for translation such as D-α-amino acids,^13^ β- and γ-amino acids,^14–17^ long-chain amino acids,^18^ aminobenzoic acid,^19^ α-aminoxy and α-hydrazino acids,^20^ α-thio acids,^21^ and others.^22–24^ The remarkable tolerance of the ribosome for such noncanonical monomers, and further expansion of its noncanonical substrate scope through engineering^25–28^ offer a possible path for repurposing mRNA-templated ribosomal translation to synthesize non-peptide polymers.

Although incorporation of structurally divergent monomers into peptides has been broadly explored *in vitro*, examples of achieving the same in living cells remain scarce. The key bottleneck limiting the progress on this front is the lack of engineered aaRSs capable of efficiently charging such monomers. In the handful of examples, where such exotic monomers were incorporated into proteins in *E. coli*, such as β^3^-*p*-Bromo-homophenylalanine^26^ and a β^2^-hydroxy acid,^29^ tRNA acylation relied on promiscuous recognition that certain wild-type aaRSs fortuitously exhibited for these substrates. The ability to systematically engineer aaRSs to charge a wider variety of noncanonical monomers with high fidelity and efficiency is needed to overcome this limitation. However, established directed evolution strategies for altering the substrate specificity of aaRSs rely on the ribosomal translation of reporter proteins with a selectable phenotype (such as antibiotic resistance, or fluorescence; Figure 1a).^30–39^ Unfortunately, such translation-dependent selection schemes are incompatible with noncanonical monomers that are poor substrates for the ribosome.

**Figure 1.**
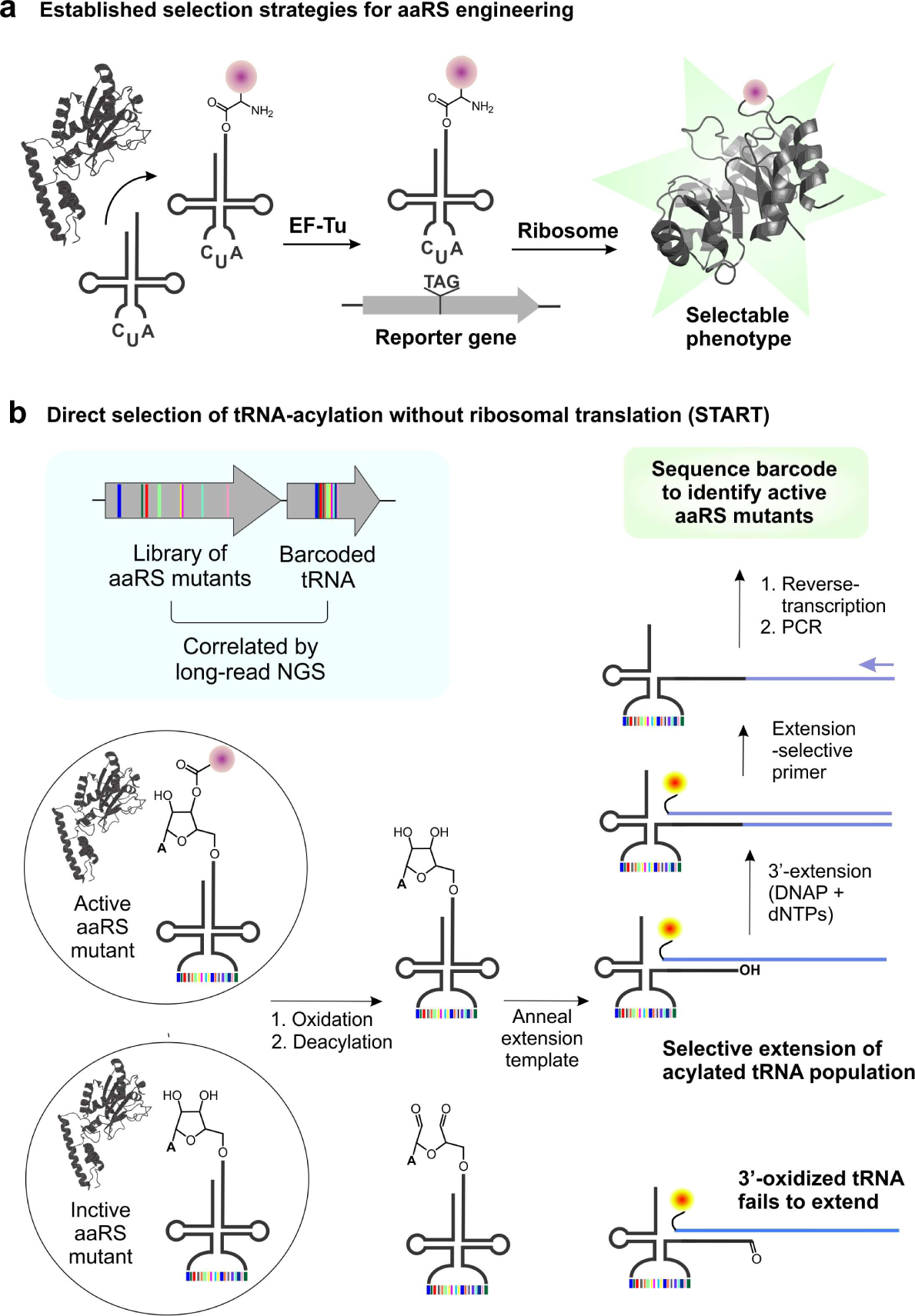
Directed evolution strategies for engineering the substrate-specificity of aaRSs. a) Established selection strategies depend on ribosomal translation to connect the activity of aaRS to the expression of a reporter protein with a selectable phenotype. This strategy is incompatible with noncanonical monomers that are poor substrates for the ribosome. b) In START, members in an aaRS mutant library are individually correlated to a sequence barcode within the cognate tRNA. Successful charging by a mutant aaRS protects its cognate barcode-containing tRNAs from a periodate-mediated oxidation, enabling their subsequent enrichment via templated 3′-extension and RT-PCR.

Here we report a general approach for engineering aaRSs that involves direct selection of tRNA acylation without ribosomal translation (START; Figure 1b). Our approach was inspired by an established strategy to enrich acylated tRNA pools from living cells, where the 2′,3′-dihydroxy functionality at the 3′-terminus of uncharged tRNAs is selectively oxidized using periodate.^40–47^ The acylated tRNAs are protected from this damage, and can be subsequently tagged with a unique oligonucleotide sequence at its intact 3′-terminus, allowing their subsequent identification through selective reverse-transcription and PCR (RT-PCR).^42–47^ Although this strategy is effective for enriching and characterizing acylated tRNA sequences, it does not reveal the identity of the aaRS responsible for their acylation. To do so, it is essential to connect the identity of each aaRS variant to its cognate tRNA sequence. In START, we achieve this connection by introducing a sequence barcode within into a permissive site within the tRNA. Using *M. alvus* pyrrolysyl-tRNA (tRNA^MaPyl^) as the model system, we demonstrate the feasibility of inserting a sequence barcode in the anticodon loop with limited impact on tRNA expression and charging. Next, the START selection scheme was optimized to allow robust enrichment of an acylated barcode-containing tRNA^MaPyl^ from a mixed population that also contained an uncharged tRNA^MaPyl^ with a distinct barcode. Then we created an active site library of *M. alvus* pyrrolysyl-tRNA synthetase (MaPylRS), where each unique mutant is encoded by associated tRNA^MaPyl^ barcodes, and subjected this library to the START scheme to identify MaPylRS mutants capable of charging different ncAAs. Finally, we show that the START selection scheme is compatible with non-α-amino acid monomers. START represents the first translation-independent directed evolution platform for aaRS engineering, which will be invaluable for developing incorporation systems for unique noncanonical monomers.

## Results and discussion

### A chemical strategy to select for charged tRNAs

A translation-independent directed evolution strategy would ideally allow the selection of active aaRS mutants simply based on their ability to acylate a cognate tRNA with a desired noncanonical monomer. Upon aaRS-catalyzed acylation of a tRNA on its 3′-terminal ribose, it becomes protected from oxidation by sodium periodate, which selectively oxidizes vicinal diol groups. This method has long been leveraged to characterize the charging status of tRNAs in cells.^40–47^ Recently, this strategy has been combined with modern enrichment and sequencing methods to perform such analyses in a more high-throughput manner.^42, 43, 45–47^ After the selective periodate-mediated oxidation of the uncharged tRNAs, a unique oligonucleotide sequence can be introduced onto the intact 3′-terminus of the acylated population either through ligation^43, 45, 46^ or by polymerase mediated extension of the 3′ terminus using a DNA template that is hybridized onto the tRNA sequence.^44^ The installed oligonucleotide sequence can be subsequently used to selectively reverse-transcribe and PCR amplify the charged tRNA sequences.

The methods described above for selective enrichment of acylated tRNAs has the potential to serve as the foundation for a translation-independent selection strategy for tRNA acylation. To explore this possibility, we focused on the *M. alvus*-derived pyrrolysyl-tRNA synthetase (MaPylRS)/tRNA^MaPyl^ pair as a model system.^48^ The pyrrolysyl pair represents the leading platform for expanding the genetic code with both ncAAs and non-α-amino acid monomers.^1–3, 9, 24, 29, 49^ We designed a 3′-fluorophore-labeled DNA template (Figure S1) that will hybridize with tRNA^MaPyl^ and allow the extension of its 3′-terminus by DNA polymerase (Figure 2a). This template selectively hybridized with tRNA^MaPyl^ expressed in *E. coli*, as observed by non-denaturing PAGE followed by fluorescence imaging; no such complex was found when total RNA from *E. coli* not expressing the tRNA^MaPyl^ was used instead (Figure 2b). Incubation of this DNA:tRNA complex with Klenow fragment of *E. coli* DNA polymerase (lacks 3′ → 5′ exonuclease activity) and dNTPs led to successful extension of the 3′-terminus, as shown by an upward shift in the PAGE analysis. Treatment of tRNA^MaPyl^ with sodium periodate prevented the appearance of this extension product, consistent with the oxidation of its free 3′-terminus (Figure S2). However, when tRNA^MaPyl^ was co-expressed with MaPylRS in the presence of a known ncAA substrate (N^ε^-Boc-L-lysine; BocK), it was protected from periodate-mediate damage and successfully yielded the extension product (Figure S2). These results confirm that the tRNA-extension strategy can be used to selectively introduce an oligonucleotide tag at the 3′-terminus of the acylated tRNA^MaPyl^ population, allowing their subsequent enrichment by RT-PCR.

**Figure 2.**
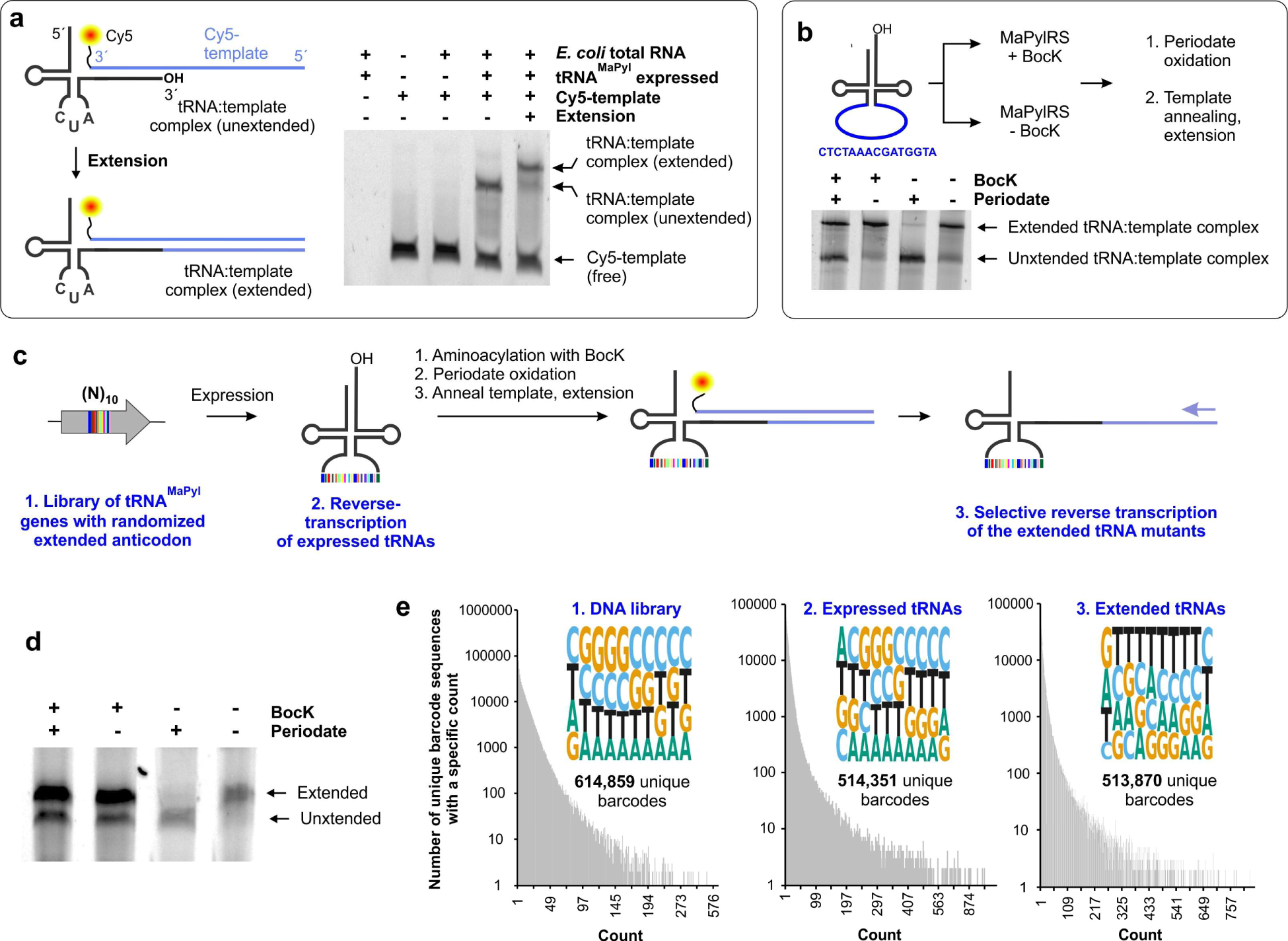
The anticodon loop of tRNA^MaPyl^ can be expanded to include a sequence barcode. a) The 3′-extension method used to selectively tag tRNA^MaPyl^ with an intact 3′-end, using a 3′-Cy5-labeled DNA template. Native PAGE followed by fluorescence imaging shows that the designed Cy5-DNA template selectively hybridizes with tRNA^MaPyl^, and enables the formation of the extension product. B) A tRNA^MaPyl^ mutant with a severely expanded anticodon loop can be effectively charged by MaPylRS as revealed by the tRNA-extension assay. In the absence of tRNA acylation, periodate treatment prevents the formation of the extension product, but when co-expressed with MaPylRS and a cognate ncAA substrate (BocK), robust formation of extension product was observed, indicating successful acylation of the extended tRNA^MaPyl^. c) Scheme for characterizing the expression and potential aminoacylation of the members of the tRNA^MaPyl^ barcode library. d) A large majority of the barcode-containing tRNA^MaPyl^ library members are successfully acylated in the presence of MaPylRS and BocK, as revealed by protection from periodate oxidation in the tRNA-extension assay. e) Illumina sequencing of the barcode-containing tRNA^MaPyl^ library DNA (left), after it is expressed in *E. coli* (middle), and members that survive periodate treatment when co-expressed with MaPylRS in the presence of BocK (right) reveal similar distribution and composition. These analyses indicate that the majority of the barcode-containing tRNA^MaPyl^ sequences are successfully expressed and acylated by MaPylRS.

### Expanding the anticodon loop of tRNA^MaPyl^ to insert a sequence barcode

To use this strategy for evolving MaPylRS, its sequence information must be encoded within tRNA^MaPyl^ such that the identity of variants responsible for tRNA-acylation can be retrieved by sequencing the acylated tRNA^MaPyl^ pool following their enrichment. We envisioned achieving this encoding by introducing a sequence barcode within a permissive site of tRNA^MaPyl^. The anticodon loop of tRNA^MaPyl^ is a promising location to introduce the barcode, given pyrrolysyl synthetases do not interact with the anticodon region.^50–55^ Indeed, the anticodon of pyrrolysyl-tRNAs have been altered to enable suppression of other nonsense codons, as well as four-base codons, indicating significant plasticity.^53–56^

To explore if the anticodon loop of tRNA^MaPyl^ can tolerate more dramatic expansions, we generated a variant where the native sequence in the anticodon loop was replaced with a random 15-nucleotide sequence (Figure 2b). The resulting expanded tRNA^MaPyl^ mutant was co-expressed in *E. coli* with MaPylRS either in the presence or absence of a cognate ncAA substrate (BocK). Total RNA isolated from these cells were subjected to tRNA-extension assay with or without periodate treatment (Figure 2b). The successful expression of the expanded tRNA^MaPyl^ mutant was confirmed by the appearance of the tRNA:DNA hybrid band. When incubated with DNA polymerase and dNTPs, the expanded tRNA^MaPyl^ successfully yielded the extension product in the absence of periodate treatment. MaPylRS was found to efficiently acylate the expanded tRNA^MaPyl^ variant with BocK, as the formation of the extension product was largely insensitive to periodate when the ncAA was included in the growth medium, but almost fully disappeared in its absence (Figure 2b).

Although this result was encouraging, we sought to further confirm that the anticodon loop of tRNA^MaPyl^ would tolerate a broad variety of sequences without significantly compromising its expression or aminoacylation by MaPylRS, which is essential to access sufficient barcode diversity for encoding a MaPylRS mutant library. To this end, we replaced the anticodon loop of tRNA^MaPyl^ with a sequence of 10 fully randomized nucleotides to create a library of roughly 10^6^ mutants (Figure 2c). Illumina sequencing (approximately 5×10^6^ usable reads) of the resulting plasmid DNA encoding this tRNA library revealed the presence of >600,000 unique barcodes (Figure 2e). Each nucleotide was well-represented at each position of the barcode library (Figure 2e). Additionally, >90% of the barcodes in the library were found to have an abundance that is within a single standard deviation from the average (∼8.5 counts/barcode), indicating reasonably good diversity and distribution of the barcode sequences. To evaluate the expression patterns of the barcode-containing tRNA^MaPyl^ genes, this plasmid library was transformed into *E. coli*, the expressed tRNA^MaPyl^ sequences were amplified by RT-PCR, and analyzed by Illumina sequencing. Over 500,000 unique sequences were identified in the expressed tRNA^MaPyl^ barcode library, and the representation of each nucleotide in the barcode was comparable to the DNA library (Figure 2e; Figure S3). These observations confirm that the majority of the barcode-containing tRNA^MaPyl^ sequences can be expressed in *E. coli*.

Finally, to explore if MaPylRS can charge barcode-containing tRNA^MaPyl^ sequences, the library was co-expressed with MaPylRS in *E. coli* in the presence of BocK. The resulting barcode-containing tRNA^MaPyl^ pool was oxidized by periodate to damage the uncharged population, and the acylated sequences subjected to tRNA-extension using a 3′-Cy5-labeled DNA template (Figure 2c). Analysis of this extension reaction by PAGE followed by fluorescence imaging (Figure 2d) showed that most of the tRNA^MaPyl^ pool was protected from periodate oxidation in the presence of BocK (but not when the ncAA was absent), indicating that a large majority of barcode-containing tRNA^MaPyl^ sequences are efficiently charged by MaPylRS. This finding was further confirmed by amplifying the acylated barcode-containing tRNA^MaPyl^ population by RT-PCR, and analyzing them by Illumina sequencing. Greater than 500,000 unique barcode sequences were identified in the acylated fraction, and their composition and distribution was similar to the library at the DNA and expressed tRNA level (Figure 2e). Together, these experiments show that expanding the tRNA^MaPyl^ anticodon loop is a viable approach for incorporating a diverse sequence barcode that does not drastically disrupt expression or charging by MaPylRS.

### Optimization of START to enrich charged, barcode-containing tRNA^MaPyl^ sequences from a mixed population

The basic steps of the START scheme involve periodate oxidation to damage the uncharged tRNAs, selective DNA-templated extension of the charged tRNAs to introduce a unique sequence at the 3′-terminus, followed by its use to amplify these sequences by RT-PCR. To improve the enrichment of the desired population, we employed a gel purification step to isolate the extended DNA:RNA hybrid from the unextended counterparts based on its different mobility on non-denaturing PAGE. We explored different lengths of the DNA template to alter the extended product size and optimize its separation from other species (Figure S4). Next, for evaluating the performance of this selection scheme, we created two standard plasmids to represent an ‘active’ or ‘inactive’ member in the library (Figure 3a). Each plasmid encoded a distinct barcode-containing tRNA^MaPyl^ sequence, but one of these also encoded wild-type MaPylRS, which will acylate the corresponding tRNA^MaPyl^ in the presence of BocK and represent an active library member. The other plasmid lacked MaPylRS and would represent an inactive library member. These plasmids were separately transformed into *E. coli*, and using the aforementioned tRNA-extension assay we confirmed that the tRNA^MaPyl^ variant co-expressed with MaPylRS was protected from oxidative damage in the presence of BocK, whereas its counterpart lacking MaPylRS was not (Figure S5). In order to quantify the degree of enrichment afforded by our strategy, the mock inactive and active tRNA^MaPyl^ populations were mixed in 10:1 ratio, and the composition of these mixed population was characterized by amplicon sequencing (∼50,000 reads) before and after subjecting it to the START scheme (Figure 3a-b). The tRNA^MaPyl^ variant co-expressed with MaPylRS was found to undergo roughly an 11-fold enrichment upon selection (Figure 3c), suggesting that this strategy should allow differentiation between barcode-containing tRNA^MaPyl^ variants associated with active or inactive MaPylRS mutants.

**Figure 3.**
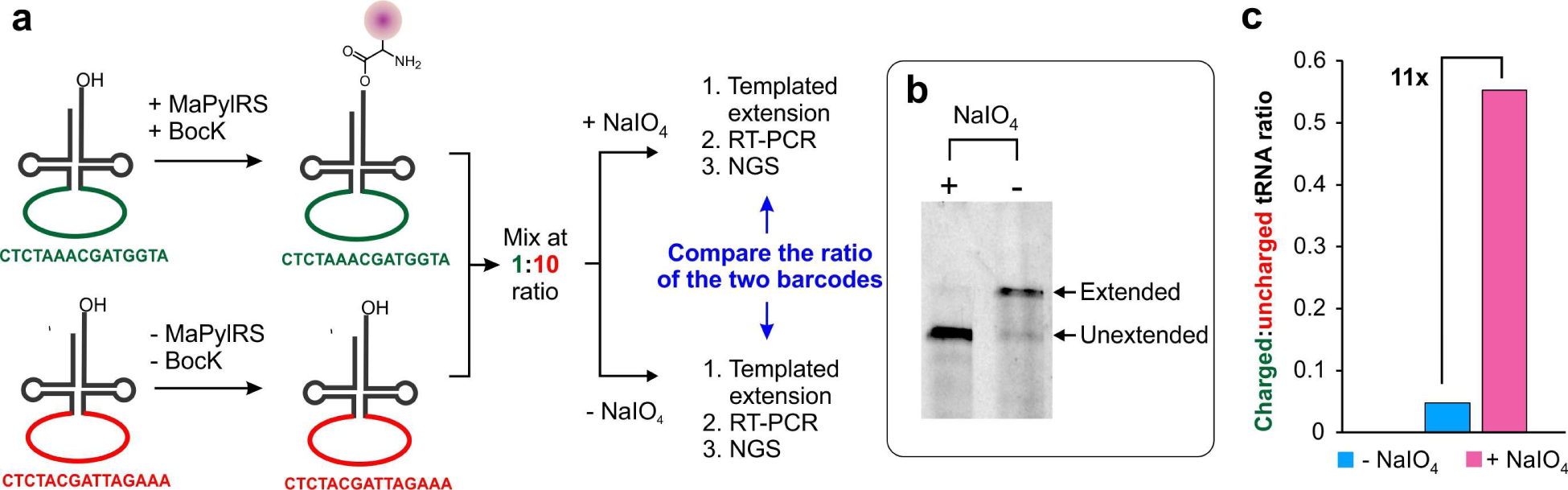
START enables enrichment of an acylated barcode-containing tRNA^MaPyl^ from a defined mixed population. a) The scheme of the experiment showing two distinct barcode-containing tRNA^MaPyl^ are separately expressed, where one is charged with BocK and the other is not. A defined mixture of these two populations is subjected to the START scheme, and their relative abundance is quantitatively measured in the presence and absence of the periodate-mediated selection using amplicon sequencing. b) PAGE followed by fluorescence imaging of the tRNA extension reaction of this mixed population in the presence or absence of periodate oxidation. c) Amplicon sequencing reveals that the acylated tRNA^MaPyl^ barcodes are enriched approximately 11-fold upon periodate selection.

### Identification of ncAA-selective MaPylRS mutants using START

Next, we explored if START can be used to identify MaPylRS mutants with altered substrate specificities from a naïve library. To this end, we simultaneously randomized three key residues (Leu125, Asn166, and Val168) around the substrate binding pocket of MaPylRS to NNK codons (Figure 4a), creating a library with a theoretical diversity of 32,768. Into the plasmid encoding the MaPylRS library, we further introduced our barcode-containing tRNA^MaPyl^ library containing an (N)_10_ randomized sequence at the anticodon loop (Figure 4b). The resulting library was covered using approximately 1.5×10^5^ transformants to ensure that A) each barcode corresponds to a distinct MaPylRS mutant, and B) each MaPylRS is associated with multiple distinct barcodes. Since this library utilizes only ∼15% of all possible barcode sequences (∼10^6^), the chances of two different MaPylRS mutants receiving the same barcode are low. Additionally, as barcode diversity in the resulting library far exceeds the diversity of the MaPylRS library (roughly 50-fold), each distinct MaPylRS mutant should be associated with multiple unique barcodes, which would be beneficial to reduce the risk of false positives.

**Figure 4.**
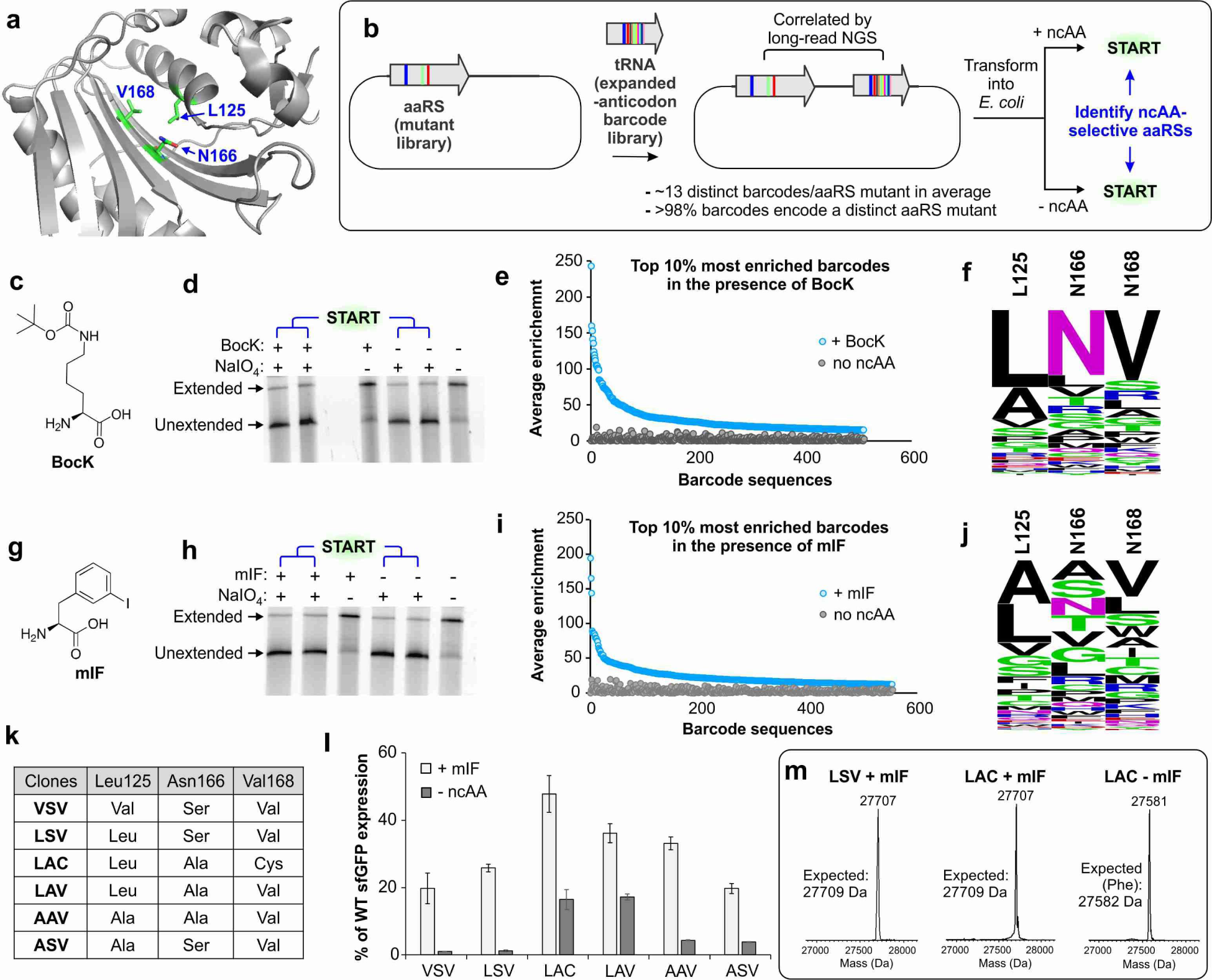
Selection of ncAA-selective MaPylRS mutants using START. a) The active site of MaPylRS, where the three key residues randomized to generate the library are highlighted. b) Scheme for generating and selecting the MaPylRS library, where each mutant is correlated to distinct barcode-containing tRNA^MaPyl^ sequences. c) The structure of BocK. d) PAGE followed by fluorescence imaging of the extension reaction on the library selection in the presence and absence of BocK. e) Top 10% most enriched sequence barcodes that show at least 2-fold higher enrichment in the presence of BocK (relative to no BocK). Enrichment of each barcode in the presence (blue) or the absence (gray) of BocK are shown. f) WebLogo analyses of the residues observed in the active sites of the MaPylRS mutants correlated to these top 10% barcodes reveal the wild-type MaPylRS sequence as the most prevalent. g) The structure of mIF. h) PAGE followed by fluorescence imaging of the extension reaction on the library selection in the presence and absence of mIF. i) Top 10% most enriched sequence barcodes that show at least 2-fold higher enrichment in the presence of mIF (relative to no mIF). Enrichment of each barcode in the presence (blue) or the absence (gray) of mIF are shown. j) WebLogo analyses of the residues observed in the active sites of the MaPylRS mutants correlated to these top 10% barcodes reveal that the prevalent signature is distinct from that of the wild-type MaPylRS. k) Selected clones for further characterizations, based on their average enrichment levels, the number of associated barcodes that show enrichment, and similarity to the consensus sequence of the most enriched sequences (panel j). l) The mIF-charging activity of each MaPylRS mutant, upon co-expression with tRNA_CUA_^MaPyl^, measured as the expression of a sfGFP-151-TAG reporter in the presence/absence of mIF and normalized to the expression of a wild-type sfGFP reporter. m) ESI-MS analysis of purified sfGFP-151-TAG reporter for LSV and LAC mutants in the presence of mIF show masses consistent with the incorporation of mIF at the TAG codon. In the absence of mIF, the LAC mutant likely charges phenylalanine, as indicated by the mass of the isolated reporter protein.

To characterize this library, and correlate each unique MaPylRS mutant to the associated sequence barcodes, we used PacBio^®^ HiFi long-read DNA sequencing. From ∼1.2×10^5^ usable reads retrieved from the sequencing analysis, we confirmed the presence of >93% of all possible MaPylRS mutants. We also found that >91% of unique MaPylRS mutants were represented by more than one barcode, with the average being ∼13 barcodes/mutant (Figure S6a). Furthermore, approximately 93% of the barcodes were found to encode a unique MaPylRS mutant (Figure S6b), which minimizes potential confusion when establishing genotype-phenotype connections.

Following characterization, the barcoded MaPylRS library was transformed into *E. coli* for subsequent selection. It is important to note that many MaPylRS active site mutants may charge one of the 20 canonical amino acids, which will render the associated barcode-containing tRNAs insensitive to periodate oxidation-mediated selection. Traditional aaRS engineering strategies typically employ a negative selection step to deplete such cross-reactive mutants.^1, 30, 31, 33, 34^ However, using the deep sequence coverage from NGS, it should be possible to characterize the enrichment pattern of nearly all mutants present in the MaPylRS library in response to the START scheme, and doing so in parallel in the presence and the absence of a desired ncAA should help identify mutants that were selectively enriched in the presence of the ncAA (Figure 4b). A similar approach was recently used for orthogonal tRNA evolution in mammalian cells.^57^

Using this strategy, the barcoded MaPylRS library was selected for its ability to charge two distinct ncAAs, BocK (Figure 4c-f) and m-iodo-L-phenylalanine (mIF) (Figure 4g-j), performed in duplicates. After selection, the surviving barcodes were amplified and characterized by Illumina sequencing. To identify barcodes associated with ncAA-selective MaPylRS mutants, we filtered the NGS data based on three key criteria: A) it must be represented in the collection of barcodes obtained by long-read sequencing characterization of the original library, B) it must show at least a 2-fold enrichment upon selection in the presence of ncAA, and C) its relative enrichment in the presence of the ncAA must be at least 2-fold higher than in the absence. Top 10% (∼500) of the surviving barcodes were arranged in a decreasing order of average enrichment in the presence of the ncAA (Figure 4e and 4i), and were used to retrieve the sequences of the corresponding MaPylRS mutants. These sequences were used to generate a WebLogo sequence map that represents the relative abundance of observed amino acid residues at each of the three randomized positions within this selected mutant pool. We were encouraged to find that the most enriched sequences for the BocK-selection matched with the wild-type MaPylRS (Figure 4f). This is expected, since BocK is a superior substrate for wild-type MaPylRS. In contrast, the most enriched mutants for the mIF-selection had a distinct sequence signature (Figure 4j). We used the following criteria to select six distinct MaPylRS mutants (Figure 4k, Figure S7) from this pool of enriched sequences for individually assessing the ability to charge mIF: A) having multiple associated barcodes that show enrichment, B) high average enrichment observed across the correlated barcodes, and C) similarity to the consensus sequence of the most enriched MaPylRS mutants. These six MaPylRS mutants were co-expressed in *E. coli* with tRNA_CUA_ ^MaPyl^, and assessed for their ability to express sfGFP-151-TAG in the presence and absence of mIF (Figure 4l). Gratifyingly, all six mutants exhibited robust reporter expression in the presence of mIF (between 20-50% wild-type sfGFP). Although four mutants (those containing VSV, LSV, AAV, and ASV at positions L125, N166, and V168, respectively) exhibited low activity in the absence of mIF, LAC and LAV mutants maintained significant (albeit significantly less than +mIF) reporter expression. ESI-MS analysis of the purified reporter protein expressed using the LSV MaPylRS mutant confirmed incorporation of only mIF, when it was supplied in the growth medium (Figure 4m). Although the MaPylRS LAC mutant showed significant reporter expression in the absence of any ncAA, it was found to selectively charge mIF when it was present during expression (Figure 4m). This behavior has been previously documented for other engineered aaRSs as well and may result from its superior affinity for the ncAA relative to the canonical counterpart.^58–60^ We also isolated the reporter protein expressed using the MaPylRS LAC variant in the absence of mIF, and observed a mass consistent with the incorporation of phenylalanine (Figure 4m). These experiments confirm that START can be used to identify novel aaRS mutants within naïve libraries that accept noncanonical monomers as substrates.

### START is compatible with non-α-amino acid monomers

Since the selection system reported here relies solely on the ability of an aaRS to acylate its cognate tRNA, it should be applicable to develop mutant aaRSs that charge noncanonical monomers whose structures diverge from that of the α-amino acids. It has been shown that the pyrrolysyl synthetases such as MaPylRS do not strongly recognize the α-amino group of the native substrate.^9^ This feature has enabled the use of this family of aaRSs to charge tRNA^Pyl^ with multiple substrate analogs in which the α-amino group is replaced with other functionalities, including α-hydroxyacids, desamino-acids, β^2^-hydroxyacids, etc., without further synthetase engineering.^10–12, 24, 26, 29^ Using this demonstrated polyspecificity of native MaPylRS, we sought to explore if the core selection system described herein is also compatible with non-α-amino acid monomers. *E. coli* cells co-expressing MaPylRS and tRNA^MaPyl^ were treated with BocK, OH-BocK, H-BocK (Figure 5a), or in the absence of a potential substrate, and the resulting tRNA population was subjected to the tRNA-extension assay with or without a periodate treatment (Figure 5b). As expected, in the absence of any potential substrate, tRNA^MaPyl^ was uncharged, and was susceptible to periodate oxidation, which prevented tRNA extension. The presence of BocK, OH-BocK, and H-BocK each facilitated the formation of the tRNA-extension product even with periodate treatment, confirming that the charging of this noncanonical monomers protects the tRNA from periodate oxidation, and that this strategy may be potentially used to identify aaRS mutants selective for structurally diverse noncanonical monomers.

**Figure 5.**
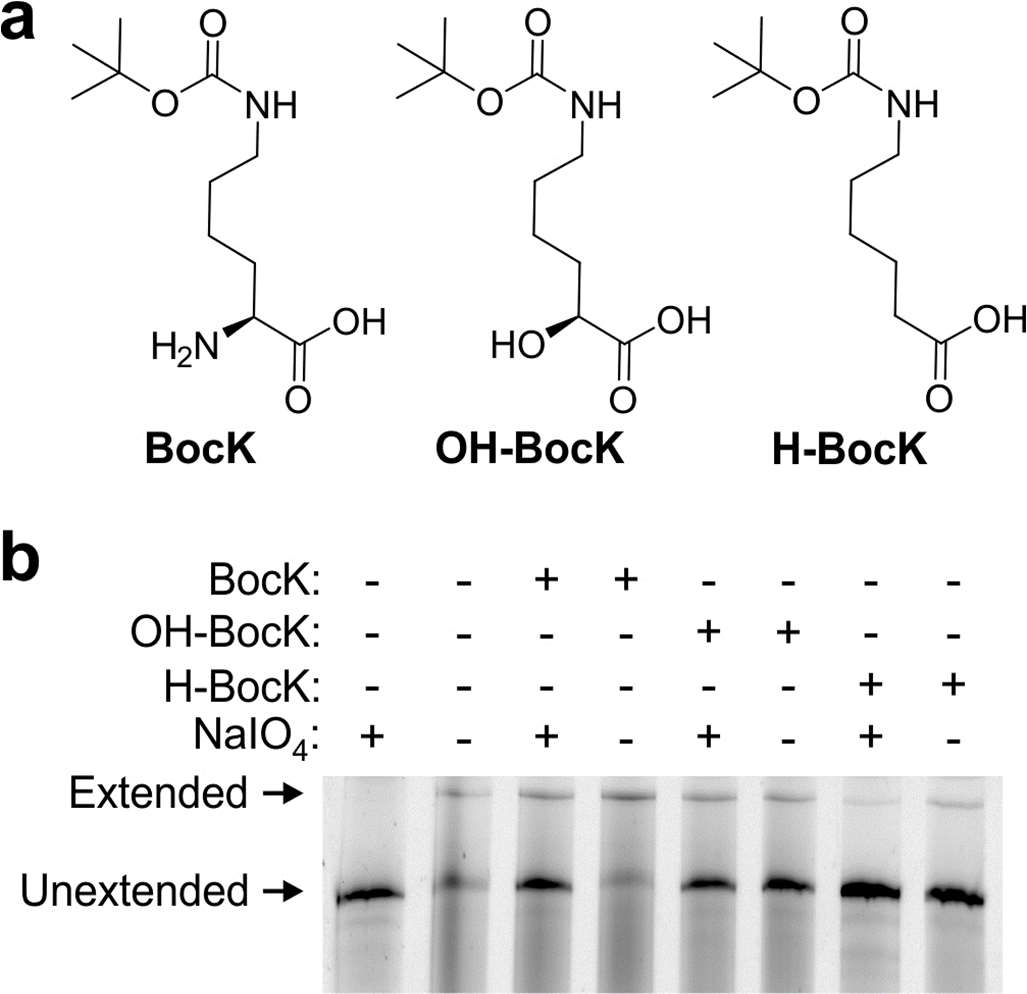
START scheme is compatible with non-α-amino acid monomers. A) The structures of BocK, OH-BocK, and H-BocK. B) Native PAGE followed by fluorescence imaging of tRNA-extension reaction performed on tRNA^MaPyl^ co-expressed with MaPylRS in the presence of BocK, OH-BocK, or H-BocK, or in the absence of any cognate substrate, and with or without periodate treatment. Successful formation of the extension product in the presence of each substrate, which are known substrates for MaPylRS, confirms the compatibility of this selection with non-α-amino acid monomers.

## Conclusion

There has been considerable interest in repurposing the mRNA-templated polypeptide synthesis by the ribosome to create novel sequence-defined polymers with noncanonical molecular architectures. However, translational incorporation of novel noncanonical monomers necessary for this purpose would require engineered variants of both aaRSs and the ribosome. Established aaRS engineering strategies are reliant on ribosomal translation, and vice versa, thereby creating an interdependence that prevents systematic introduction of structurally novel noncanonical monomers into the genetic code. We offer a solution to this challenge by developing the first directed evolution strategy for engineering aaRSs, START, which does not rely on ribosomal translation. We show that tRNAs can be equipped with sequence barcodes, which can be correlated to individual mutants in large aaRS libraries through long-read next-generation sequencing. Acylation of the correlated partner tRNA by a novel aaRS mutant protects it from periodate-mediated oxidation, allowing their subsequent enrichment. The identity of the corresponding aaRS mutant can then be retrieved from the barcode sequence. The efficacy of this strategy was demonstrated by identifying novel mutants of MaPylRS selective for two ncAAs from a naïve library. The compatibility of this selection system with non-α-amino acids was further demonstrated using the polyspecificity of MaPylRS for such substrates. It should be possible to extend the strategy described here to other aaRS/tRNA pairs. Although many aaRSs recognize the anticodon of its cognate tRNA as an identity element, which may complicate the introduction of a barcode in this region, it has been possible engineer aaRS anticodon binding domains to be permissive for non-natural expanded anticodon sequences.^61, 62^ Furthermore, such sequence barcodes can also be inserted into alternative locations within the tRNA to avoid aaRS identity elements. Indeed, it has been shown that aaRSs can effectively acylate significantly altered versions of their cognate tRNAs, including severely miniaturized versions.^63–66^ In summary, START offers an exciting opportunity to develop aaRS mutants to charge structurally divergent noncanonical monomers, and enable the ribosomal synthesis of evolvable, sequence-defined polymers with unprecedented structure and function.

## ASSOCIATED CONTENT

### Supporting Information

The Supporting Information is available free of charge on the ACS Publications website. Experimental methods, nucleotide sequences, supplementary figures and tables. (PDF) **AUTHOR INFORMATION**

### Notes

AC and CS are co-inventors on a patent application that incorporates methods outlined in this manuscript. AC is a co-founder and senior advisor at BrickBio, Inc.

## ACKNOWLEDGMENT

This work was supported by the NSF Center for Genetically Encoded Materials (C-GEM; CHE 2002182). We thank Huiqing Zhou and Mitch Tepe (Boston College) for helpful discussions and assistance with the RNA workflow.

## References

1. Chin, J. W. (2017) Expanding and reprogramming the genetic code, Nature 550, 53–60.

2. Dumas, A., Lercher, L., Spicer, C. D., and Davis, B. G. (2015) Designing logical codon reassignment - Expanding the chemistry in biology, Chemical science 6, 50–69.

3. Young, D. D., and Schultz, P. G. (2018) Playing with the molecules of life, ACS chemical biology.

4. Icking, L.-S., Riedlberger, A. M., Krause, F., Widder, J., Frederiksen, Anne S., Stockert, F., Spädt, M., Edel, N., Armbruster, D., Forlani, G., Franchini, S., Kaas, P., Kırpat Konak, B. M., Krier, F., Lefebvre, M., Mazraeh, D., Ranniger, J., Gerstenecker, J., Gescher, P., Voigt, K., Salavei, P., Gensch, N., Di Ventura, B., and Öztürk, M. A. (2023) iNClusive: a database collecting useful information on non-canonical amino acids and their incorporation into proteins for easier genetic code expansion implementation, Nucleic Acids Research.

5. Vargas-Rodriguez, O., Sevostyanova, A., Söll, D., and Crnković, A. (2018) Upgrading aminoacyl-tRNA synthetases for genetic code expansion, Current opinion in chemical biology 46, 115–122.

6. Manandhar, M., Chun, E., and Romesberg, F. E. (2021) Genetic Code Expansion: Inception, Development, Commercialization, Journal of the American Chemical Society 143, 4859–4878.

7. Sun, S. B., Schultz, P. G., and Kim, C. H. (2014) Therapeutic applications of an expanded genetic code, Chembiochem : a European journal of chemical biology 15, 1721–1729.

8. Guo, J., Wang, J., Anderson, J. C., and Schultz, P. G. (2008) Addition of an α-Hydroxy Acid to the Genetic Code of Bacteria, Angewandte Chemie International Edition 47, 722–725.

9. Kobayashi, T., Yanagisawa, T., Sakamoto, K., and Yokoyama, S. (2009) Recognition of non-alpha-amino substrates by pyrrolysyl-tRNA synthetase, Journal of molecular biology 385, 1352–1360.

10. Li, Y. M., Yang, M. Y., Huang, Y. C., Li, Y. T., Chen, P. R., and Liu, L. (2012) Ligation of expressed protein α-hydrazides via genetic incorporation of an α-hydroxy acid, ACS chemical biology 7, 1015–1022.

11. Bindman, N. A., Bobeica, S. C., Liu, W. R., and van der Donk, W. A. (2015) Facile Removal of Leader Peptides from Lanthipeptides by Incorporation of a Hydroxy Acid, J Am Chem Soc 137, 6975–6978.

12. Spinck, M., Piedrafita, C., Robertson, W. E., Elliott, T. S., Cervettini, D., de la Torre, D., and Chin, J. W. (2023) Genetically programmed cell-based synthesis of non-natural peptide and depsipeptide macrocycles, Nature chemistry 15, 61–69.

13. Katoh, T., Iwane, Y., and Suga, H. (2017) Logical engineering of D-arm and T-stem of tRNA that enhances d-amino acid incorporation, Nucleic Acids Res 45, 12601–12610.

14. Katoh, T., and Suga, H. (2020) Ribosomal Elongation of Cyclic γ-Amino Acids using a Reprogrammed Genetic Code, J Am Chem Soc 142, 4965–4969.

15. Adaligil, E., Song, A., Cunningham, C. N., and Fairbrother, W. J. (2021) Ribosomal Synthesis of Macrocyclic Peptides with Linear γ(4)- and β-Hydroxy-γ(4)-amino Acids, ACS chemical biology 16, 1325–1331.

16. Fujino, T., Goto, Y., Suga, H., and Murakami, H. (2016) Ribosomal Synthesis of Peptides with Multiple β-Amino Acids, J Am Chem Soc 138, 1962–1969.

17. Sando, S., Abe, K., Sato, N., Shibata, T., Mizusawa, K., and Aoyama, Y. (2007) Unexpected preference of the E. coli translation system for the ester bond during incorporation of backbone-elongated substrates, J Am Chem Soc 129, 6180–6186.

18. Lee, J., Schwarz, K. J., Kim, D. S., Moore, J. S., and Jewett, M. C. (2020) Ribosome-mediated polymerization of long chain carbon and cyclic amino acids into peptides in vitro, Nature communications 11, 4304.

19. Katoh, T., and Suga, H. (2020) Ribosomal Elongation of Aminobenzoic Acid Derivatives, J Am Chem Soc 142, 16518–16522.

20. Katoh, T., and Suga, H. (2021) Consecutive Ribosomal Incorporation of α-Aminoxy/α-Hydrazino Acids with l/d-Configurations into Nascent Peptide Chains, J Am Chem Soc 143, 18844–18848.

21. Takatsuji, R., Shinbara, K., Katoh, T., Goto, Y., Passioura, T., Yajima, R., Komatsu, Y., and Suga, H. (2019) Ribosomal Synthesis of Backbone-Cyclic Peptides Compatible with In Vitro Display, J Am Chem Soc 141, 2279–2287.

22. Ad, O., Hoffman, K. S., Cairns, A. G., Featherston, A. L., Miller, S. J., Söll, D., and Schepartz, A. (2019) Translation of Diverse Aramid- and 1,3-Dicarbonyl-peptides by Wild Type Ribosomes in Vitro, ACS central science 5, 1289–1294.

23. Lee, J., Coronado, J. N., Cho, N., Lim, J., Hosford, B. M., Seo, S., Kim, D. S., Kofman, C., Moore, J. S., Ellington, A. D., Anslyn, E. V., and Jewett, M. C. (2022) Ribosome-mediated biosynthesis of pyridazinone oligomers in vitro, Nature communications 13, 6322.

24. Fricke, R., Swenson, C. V., Roe, L. T., Hamlish, N. X., Shah, B., Zhang, Z., Ficaretta, E., Ad, O., Smaga, S., Gee, C. L., Chatterjee, A., and Schepartz, A. (2023) Expanding the substrate scope of pyrrolysyl-transfer RNA synthetase enzymes to include non-α-amino acids in vitro and in vivo, Nature chemistry 15, 960–971.

25. Dedkova, L. M., Fahmi, N. E., Golovine, S. Y., and Hecht, S. M. (2003) Enhanced D-amino acid incorporation into protein by modified ribosomes, J Am Chem Soc 125, 6616–6617.

26. Melo Czekster, C., Robertson, W. E., Walker, A. S., Söll, D., and Schepartz, A. (2016) In Vivo Biosynthesis of a β-Amino Acid-Containing Protein, J Am Chem Soc 138, 5194–5197.

27. Maini, R., Dedkova, L. M., Paul, R., Madathil, M. M., Chowdhury, S. R., Chen, S., and Hecht, S. M. (2015) Ribosome-Mediated Incorporation of Dipeptides and Dipeptide Analogues into Proteins in Vitro, Journal of the American Chemical Society 137, 11206–11209.

28. Chen, S., Ji, X., Gao, M., Dedkova, L. M., and Hecht, S. M. (2019) In Cellulo Synthesis of Proteins Containing a Fluorescent Oxazole Amino Acid, Journal of the American Chemical Society 141, 5597–5601.

29. Hamlish, N. X., Abramyan, A. M., and Schepartz, A. (2023) Incorporation of multiple β2-backbones into a protein in vivo using an orthogonal aminoacyl-tRNA synthetase, bioRxiv, 2023.2011.2007.565973.

30. Liu, D. R., and Schultz, P. G. (1999) Progress toward the evolution of an organism with an expanded genetic code, Proceedings of the National Academy of Sciences of the United States of America 96, 4780–4785.

31. Wang, L., Brock, A., Herberich, B., and Schultz, P. G. (2001) Expanding the genetic code of Escherichia coli, Science (New York, N.Y.) 292, 498–500.

32. Chin, J. W., Cropp, T. A., Anderson, J. C., Mukherji, M., Zhang, Z., and Schultz, P. G. (2003) An expanded eukaryotic genetic code, Science (New York, N.Y.) 301, 964–967.

33. Santoro, S. W., Wang, L., Herberich, B., King, D. S., and Schultz, P. G. (2002) An efficient system for the evolution of aminoacyl-tRNA synthetase specificity, Nature Biotechnology 20, 1044–1048.

34. Ellefson, J. W., Meyer, A. J., Hughes, R. A., Cannon, J. R., Brodbelt, J. S., and Ellington, A. D. (2014) Directed evolution of genetic parts and circuits by compartmentalized partnered replication, Nat Biotechnol 32, 97–101.

35. Bryson, D. I., Fan, C., Guo, L. T., Miller, C., Söll, D., and Liu, D. R. (2017) Continuous directed evolution of aminoacyl-tRNA synthetases, Nature chemical biology 13, 1253–1260.

36. Suzuki, T., Miller, C., Guo, L. T., Ho, J. M. L., Bryson, D. I., Wang, Y. S., Liu, D. R., and Söll, D. (2017) Crystal structures reveal an elusive functional domain of pyrrolysyl-tRNA synthetase, Nature chemical biology 13, 1261–1266.

37. Fischer, J. T., Söll, D., and Tharp, J. M. (2022) Directed Evolution of Methanomethylophilus alvus Pyrrolysyl-tRNA Synthetase Generates a Hyperactive and Highly Selective Variant, Frontiers in molecular biosciences 9, 850613.

38. Stieglitz, J. T., and Van Deventer, J. A. (2022) High-Throughput Aminoacyl-tRNA Synthetase Engineering for Genetic Code Expansion in Yeast, ACS Synthetic Biology 11, 2284–2299.

39. Hohl, A., Karan, R., Akal, A., Renn, D., Liu, X., Ghorpade, S., Groll, M., Rueping, M., and Eppinger, J. (2019) Engineering a Polyspecific Pyrrolysyl-tRNA Synthetase by a High Throughput FACS Screen, Scientific reports 9, 11971.

40. Dyer, J. R. (1956) Use of periodate oxidations in biochemical analysis, Methods of biochemical analysis 3, 111–152.

41. Dittmar, K. A., Sørensen, M. A., Elf, J., Ehrenberg, M., and Pan, T. (2005) Selective charging of tRNA isoacceptors induced by amino-acid starvation, EMBO reports 6, 151–157.

42. Evans, M. E., Clark, W. C., Zheng, G., and Pan, T. (2017) Determination of tRNA aminoacylation levels by high-throughput sequencing, Nucleic Acids Research 45, e133–e133.

43. Erber, L., Hoffmann, A., Fallmann, J., Betat, H., Stadler, P. F., and Mörl, M. (2020) LOTTE-seq (Long hairpin oligonucleotide based tRNA high-throughput sequencing): specific selection of tRNAs with 3’-CCA end for high-throughput sequencing, RNA biology 17, 23–32.

44. Cervettini, D., Tang, S., Fried, S. D., Willis, J. C. W., Funke, L. F. H., Colwell, L. J., and Chin, J. W. (2020) Rapid discovery and evolution of orthogonal aminoacyl-tRNA synthetase–tRNA pairs, Nature Biotechnology 38, 989–999.

45. Passarelli, M. C., Pinzaru, A. M., Asgharian, H., Liberti, M. V., Heissel, S., Molina, H., Goodarzi, H., and Tavazoie, S. F. (2022) Leucyl-tRNA synthetase is a tumour suppressor in breast cancer and regulates codon-dependent translation dynamics, Nature Cell Biology 24, 307–315.

46. Davidsen, K., and Sullivan, L. B. (2023) A robust method for measuring aminoacylation through tRNA-Seq, eLife Sciences Publications, Ltd.

47. Tsukamoto, Y., Nakamura, Y., Hirata, M., Sakate, R., and Kimura, T. (2022) i-tRAP (individual tRNA acylation PCR): A convenient method for selective quantification of tRNA charging, RNA (New York, N.Y.) 29, 111–122.

48. Willis, J. C. W., and Chin, J. W. (2018) Mutually orthogonal pyrrolysyl-tRNA synthetase/tRNA pairs, Nature chemistry 10, 831–837.

49. Wan, W., Tharp, J. M., and Liu, W. R. (2014) Pyrrolysyl-tRNA synthetase: an ordinary enzyme but an outstanding genetic code expansion tool, Biochimica et biophysica acta 1844, 1059–1070.

50. Nozawa, K., O’Donoghue, P., Gundllapalli, S., Araiso, Y., Ishitani, R., Umehara, T., Söll, D., and Nureki, O. (2009) Pyrrolysyl-tRNA synthetase-tRNA(Pyl) structure reveals the molecular basis of orthogonality, Nature 457, 1163–1167.

51. Seki, E., Yanagisawa, T., Kuratani, M., Sakamoto, K., and Yokoyama, S. (2020) Fully Productive Cell-Free Genetic Code Expansion by Structure-Based Engineering of Methanomethylophilus alvus Pyrrolysyl-tRNA Synthetase, ACS Synthetic Biology 9, 718–732.

52. Ambrogelly, A., Gundllapalli, S., Herring, S., Polycarpo, C., Frauer, C., and Söll, D. (2007) Pyrrolysine is not hardwired for cotranslational insertion at UAG codons, Proceedings of the National Academy of Sciences of the United States of America 104, 3141–3146.

53. Chatterjee, A., Sun, S. B., Furman, J. L., Xiao, H., and Schultz, P. G. (2013) A Versatile Platform for Single- and Multiple-Unnatural Amino Acid Mutagenesis in Escherichia coli, Biochemistry 52, 1828–1837.

54. Zheng, Y., Addy, P. S., Mukherjee, R., and Chatterjee, A. (2017) Defining the current scope and limitations of dual noncanonical amino acid mutagenesis in mammalian cells, Chemical science 8, 7211–7217.

55. Niu, W., Schultz, P. G., and Guo, J. (2013) An expanded genetic code in mammalian cells with a functional quadruplet codon, ACS chemical biology 8, 1640–1645.

56. Dunkelmann, D. L., Willis, J. C. W., Beattie, A. T., and Chin, J. W. (2020) Engineered triply orthogonal pyrrolysyl-tRNA synthetase/tRNA pairs enable the genetic encoding of three distinct non-canonical amino acids, Nature chemistry 12, 535–544.

57. Jewel, D., Kelemen, R. E., Huang, R. L., Zhu, Z., Sundaresh, B., Cao, X., Malley, K., Huang, Z., Pasha, M., Anthony, J., van Opijnen, T., and Chatterjee, A. (2023) Virus-assisted directed evolution of enhanced suppressor tRNAs in mammalian cells, Nature methods 20, 95–103.

58. Young, D. D., Young, T. S., Jahnz, M., Ahmad, I., Spraggon, G., and Schultz, P. G. (2011) An evolved aminoacyl-tRNA synthetase with atypical polysubstrate specificity, Biochemistry 50, 1894–1900.

59. Italia, J. S., Addy, P. S., Wrobel, C. J. J., Crawford, L. A., Lajoie, M. J., Zheng, Y., and Chatterjee, A. (2017) An orthogonalized platform for genetic code expansion in both bacteria and eukaryotes, Nature chemical biology 13, 446–450.

60. Italia, J. S., Latour, C., Wrobel, C. J. J., and Chatterjee, A. (2018) Resurrecting the Bacterial Tyrosyl-tRNA Synthetase/tRNA Pair for Expanding the Genetic Code of Both E. coli and Eukaryotes, Cell chemical biology 25, 1304–1312.e1305.

61. Chatterjee, A., Xiao, H., and Schultz, P. G. (2012) Evolution of multiple, mutually orthogonal prolyl-tRNA synthetase/tRNA pairs for unnatural amino acid mutagenesis in Escherichia coli, Proceedings of the National Academy of Sciences of the United States of America 109, 14841–14846.

62. Neumann, H., Slusarczyk, A. L., and Chin, J. W. (2010) De Novo Generation of Mutually Orthogonal Aminoacyl-tRNA Synthetase/tRNA Pairs, Journal of the American Chemical Society 132, 2142–2144.

63. Buechter, D. D., Schimmel, P., and de Duve, C. (1993) Aminoacylation of RNA Minihelices: Implications for tRNA Synthetase Structural Design and Evolution, Critical Reviews in Biochemistry and Molecular Biology 28, 309–322.

64. Frugier, M., Florentz, C., and Giegé, R. (1994) Efficient aminoacylation of resected RNA helices by class II aspartyl-tRNA synthetase dependent on a single nucleotide, The EMBO journal 13, 2218–2226.

65. Shiba, K., Ripmaster, T., Suzuki, N., Nichols, R., Plotz, P., Noda, T., and Schimmel, P. (1995) Human alanyl-tRNA synthetase: conservation in evolution of catalytic core and microhelix recognition, Biochemistry 34, 10340–10349.

66. Martinis, S. A., and Schimmel, P. (1994) Small RNA Oligonucleotide Substrates for Specific Aminoacylations, In tRNA, pp 349–370.

