## Supporting information for "A translation-independent directed evolution strategy to engineer aminoacyl-tRNA synthetases"

**Supporting information figures**:

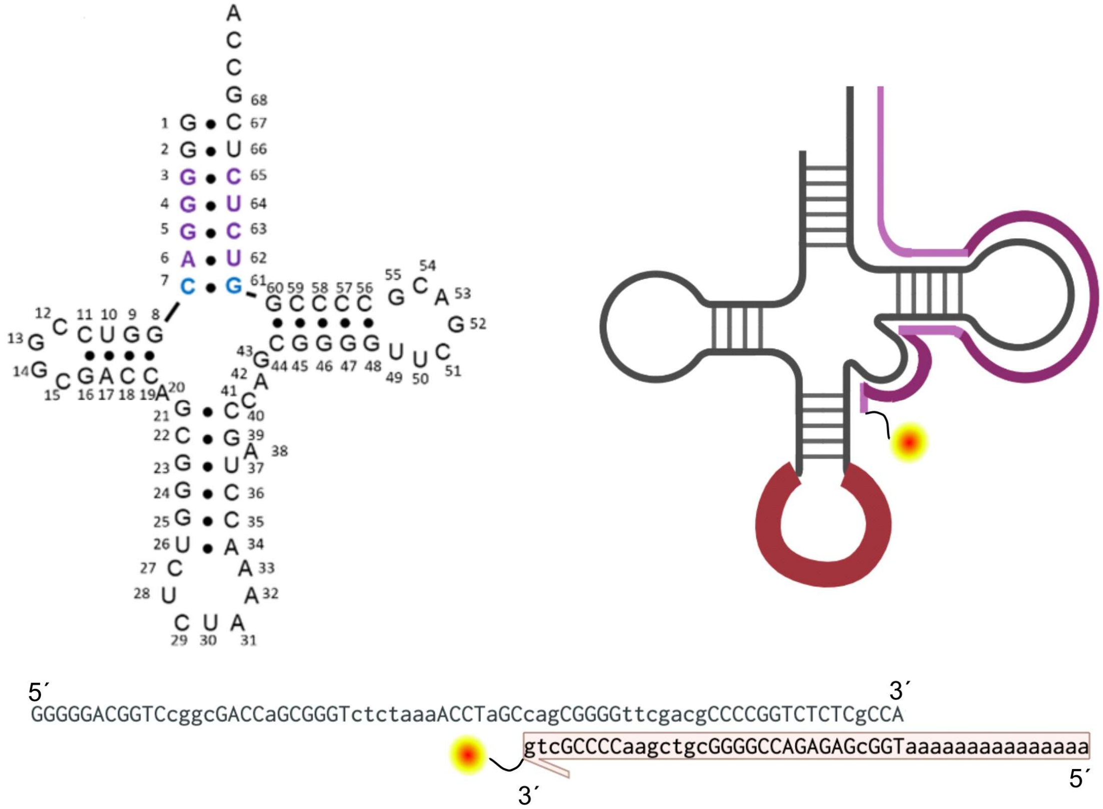


**Figure S1**. The design of a 3′-Cy5-labeled DNA template for selective 3′-extension of tRNA$^{MaPyl}$

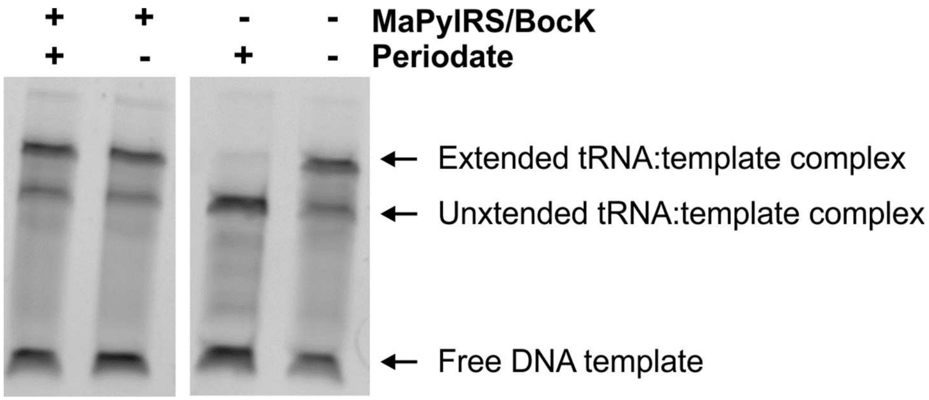


**Figure S2**. The tRNA-3'-extension strategy differentiates charged and uncharged $tRNA^{MaPyl}$. When $tRNA^{MaPyl}$ is co-expressed with MaPylRS in the presence of BocK, it is protected from periodate oxidation and can form the extension product. However, in the absence of MaPylRS, the uncharged tRNA is oxidized by periodate and is unable to form an extension product.

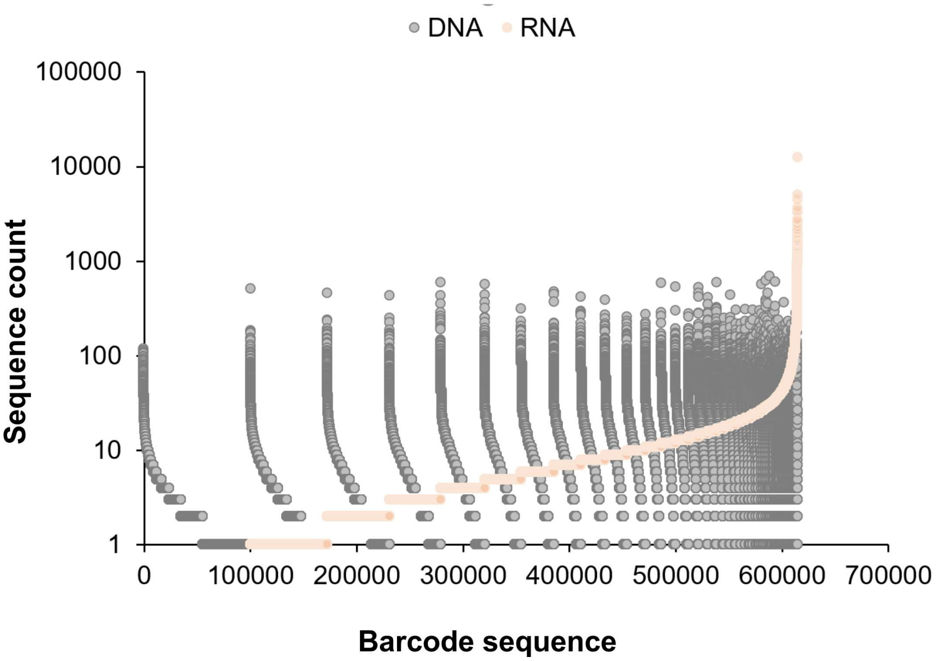


**Figure S3**. The majority of the barcodes observed in the tRNA$^{MaPyl}$ gene library are also found when the expressed tRNAs are sequenced following RT-PCR. Barcodes are organized by their observed abundance within the expressed tRNA pool (orange dots), and the corresponding abundance within the gene pool (gray dots) are co-plotted. Out of the >600,000 sequenced barcodes observed in the gene pool, >500,000 are also found within the expressed tRNA pool.

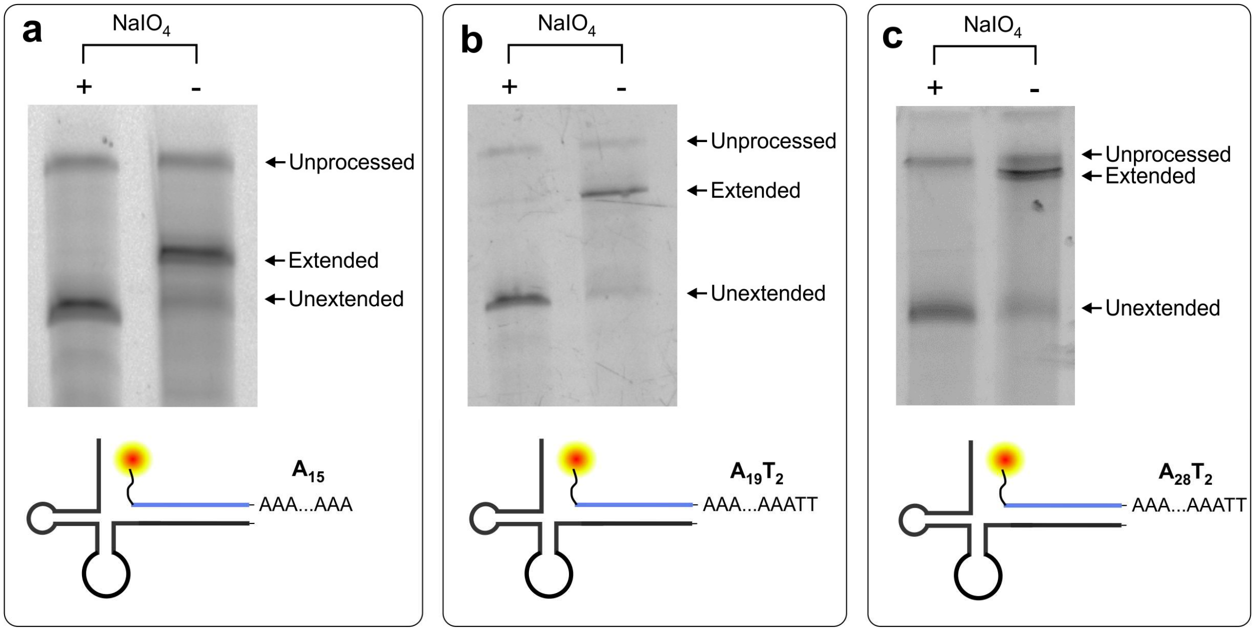


**Figure S4**. Optimization of the DNA template length for efficient separation of the tRNA-extension product from other $tRNA^{MaPyl}$ species using PAGE (lower molecular weight unextended counterpart, and the higher molecular weight band, which is likely a pre-tRNA that was not appropriately processed to generate mature tRNA).

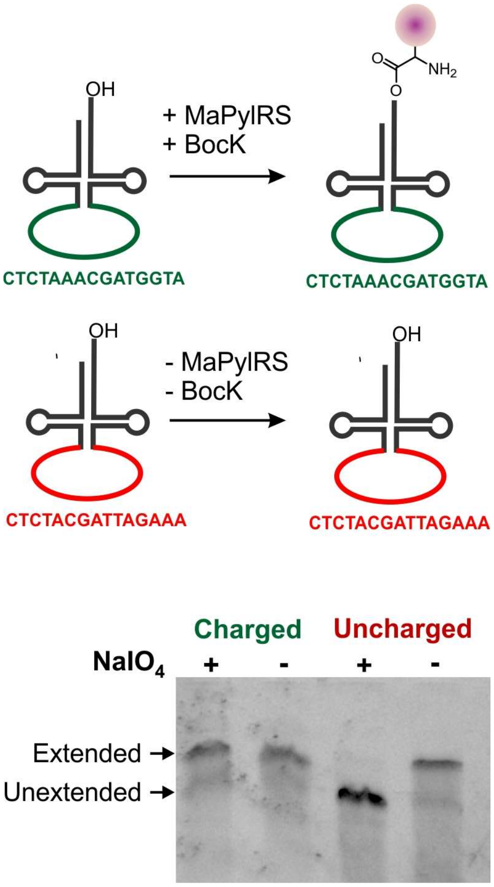


**Figure S5**. Two distinct barcode-containing $tRNA^{MaPyl}$ sequences used to create a defined mixture of charged and uncharged species. One of these was co-expressed with MaPylRS in the presence of BocK, while the other was expressed in the absence of its cognate synthetase and BocK. The tRNA-extension assay shows that the charged $tRNA^{MaPyl}$ is protected from periodate oxidation, as shown by the formation of the extended product. Whereas the extension of the uncharged tRNA is sensitive to periodate treatment.

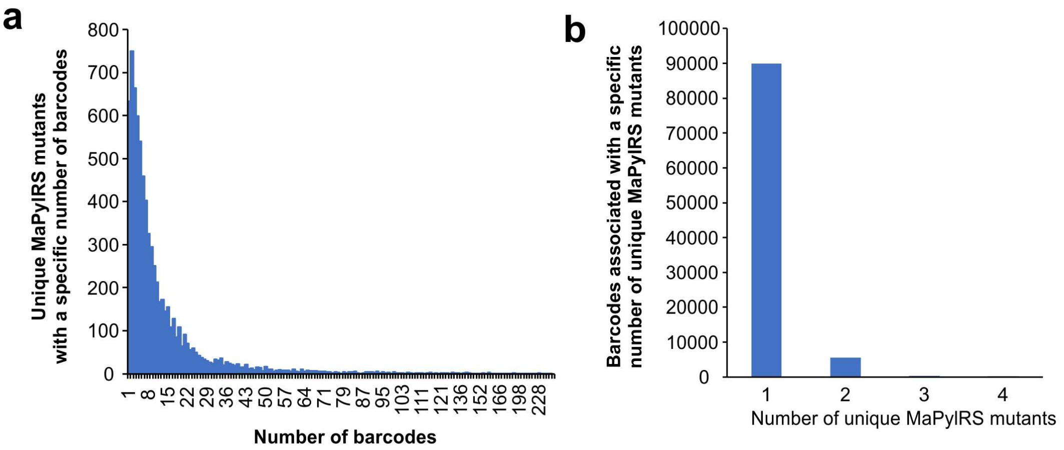


**Figure S6**. Long-read PacBio sequencing of the tRNA-barcoded MaPylRS library reveals that: a) the majority of the unique MaPylRS mutants are associated with more than one distinct barcode, with the average being ~13 barcodes/MaPylRS mutant (note: the wild-type $tRNA^{MaPyl}$ sequence was removed from this analysis due to its overabundance), and b) nearly all barcodes are correlated to a unique MaPylRS.

| L125/N166/V168 | Similarity to consensus | Number of barcodes in top 10% (>2, 2, <2) | Average enrichment across all barcodes (ranked based on percentile, >95th, >90th, <90th) |
|---|---|---|---|
| LAV | **High** | 3 | 73.4 |
| AAV | **High** | 2 | 48.8 |
| ASV | **High** | 5 | 21.4 |
| VSV | **High** | 2 | 50.7 |
| LSV | **High** | 5 | 26.8 |
| LAC | **Medium** | 1 | 194.2 |

**Figure S7**. Criteria used to select promising individual MaPylRS mutants from the selected pool for further characterization

**Material and Methods:**

**Safety considerations**

No unexpected or significant hazards were anticipated or encountered.

**Noncanonical Amino Acids**

$N^{\varepsilon}$-Boc-L-Lysine was purchased from Chem-Impex International. 2-Amino-3-(3-iodophenyl)propanoic acid (m-iodo-L-phenylalanine) was purchased from Ambeed Inc. Boc-6-aminohexanoic acid (H-BocK) was purchased from Sigma-Aldrich. OH-BocK was a gift from Prof. Alanna Schepartz.

**General:**

For all cloning, the *E. coli* DH10B electrocompetent strain was used for transformation as well as plasmid propagation, and the cells were cultured using Luria broth (LB) for solid and liquid media. All PCRs were carried out using Phusion Hot Start II DNA Polymerase (Thermo Scientific) according to the manufacturer's instructions. Restriction enzymes and T4 DNA ligase were purchased from New England Biolabs. Gibson assembly HiFi mix was purchased from Fisher Scientific. All custom DNA oligos including randomized primers and tREX probes were purchased from Integrated DNA Technologies (IDT). All other oligos were purchased from GENEWIZ. Sanger sequencing was performed by GENEWIZ. Amplicon sequencing was performed by Quintara Biosciences. Whole plasmid sequencing was performed by Plasmidsaurus. Long-read PacBio sequencing was performed by Quintara Biosciences.

**Construction of extended anticodon tRNA mutants:**

| **Extended tRNA anticodons** | **Sequence (5' to 3')** |
|---|---|
| 15AC1 | CTCTAAACGATGGTA |
| 15AC2 | CTCTACGATTAGAAA |

The two extended anticodon tRNAs were cloned into pBK lpp-$tRNA^{MaPyl}$ and pBK lpp-$tRNA^{MaPyl}$-metG-MaPylRS plasmids using overlap extension cloning. The two overlapping fragments were generated with lpp_BamHI_F + MaPylT_15AC_R, MaPylT_15AC_F + tRNA_NotI_R. The two fragments were joined together with an overlap extension PCR with terminal primers lpp_BamHI_F and tRNA_NotI_R. The amplified product was purified with 1% agarose gel and digested with BamHI/NotI enzymes, followed by ligation with T4 DNA ligase into the pBK vector to generate pBK lpp- $tRNA^{MaPyl}$-15AC1 and pBK lpp-$tRNA^{MaPyl}$-15AC1-metG-MaPylRS. The insertion of the extended tRNA was confirmed via Sanger sequencing.

For extended anticodon 15AC2, the two fragments were generated using lpp_BamHI_F + MaPyl_15AC2_iR, MaPyl_15AC2_iF + tRNA_NotI_R. The constructs were cloned as described above.

**Construction of 10-nucleotide (nt) extended anticodon tRNA library:**

The 10-nt extended anticodon tRNA library (1,048,576 or ~$10^6$ variants) was constructed using overlap extension similarly as described above. The primers used to generate the overlapping fragments were lpp_BamHI_F + MaPyl_lib_R, MaPyl_lib_10N_F + tRNA_NotI_R. The amplified insert was purified with 1% agarose gel, digested with BamHI/NotI and ligated into 5 µg of pBK lpp-$tRNA^{MaPyl}$-15AC1-metG-MaPylRS vector, replacing the $tRNA^{MaPyl}$ 15AC1. The ligation product was ethanol precipitated overnight with DNA: 3M NaOAc (pH 5.2): yeast tRNA (10 mg/mL): 100% cold ethanol in 1: 0.1: 0.008: 3 parts. The mixture was spun down at 20,000 xg for 20 mins at 4°C, washed with 70% ethanol, spun down again for 10 mins. The supernatant was dumped, the pellet was air-dried for 15 mins and finally, resuspended in 10 µL of $ddH_2O$. The ethanol precipitated DNA was transformed into 500 µL DH10B electrocompetent cells, recovered for 2 hours, spun down, resuspended in 1 mL LB, and plated on 15 cm dishes. The library was covered by ~$5*10^9$ distinct cfus (~5000-fold coverage). These colonies were pooled and stored as glycerol stocks for further analyses.

**Construction of tRNA barcoded-MaPylRS library:**

To construct the tRNA barcoded-MaPylRS library, first the MaPylRS library (32,768 variants) was generated using overlap extension. Three different fragments A, B and C were generated with the following set of primers: metG_NotI_F + MaPylRS_123_iR, MaPylRS_L125NNK_iF + MaPylRS_L165_iR and MaPylRS_N166, V168 NNK_iF + MaPylRS_NcoI_R. The pieces A and B were joined together using a primerless-overlap extension PCR. Subsequently, another primerless overlap extension PCR was performed to join the piece AB with piece C. The amplified insert was purified with 1% agarose gel, digested with NotI/NcoI and ligated into 1.3 µg of pBK lpp-$tRNA^{MaPyl}$-15AC1-metG-MaPylRS. The ligation product was ethanol precipitated as described above and later transformed into 250 µL DH10B electrocompetent cells, recovered for 2 hours, spun down, resuspended in 1 mL LB, and plated on 15 cm dishes. The library was covered with ~$4*10^7$ distinct cfus (~1000-fold coverage). The diversity of the sequencing was analyzed by sequencing 20 different clones as well as Amplicon sequencing.

After characterizing the MaPylRS library, the 10-nt extended anticodon tRNA library was cloned into the pBK vector containing MaPylRS library as described previously. The 10-nt extended anticodon tRNA library was capped at 50-fold coverage (~1.6 * $10^6$ variants) to minimize the number of barcodes corresponding to multiple MaPylRS mutants.

**NGS Analysis for input library:**

NGS Analysis for long-read PacBio sequencing was performed with the following script: https://github.com/cpsoni74/PacBio_seq_analysis/tree/d5e50606fd13cb241a1ec8d4cd27bf5c670498b1

The reads were first aligned to the template using the mapping tool minimap2 and the reads that misaligned were discarded. The reads were then filtered based on data quality using Phred score (Q-score) as the metric. Only reads in which each base of the tRNA barcode had a Q-score >30 were considered. Subsequently, all the MaPylRS sequences corresponding to these reads were extracted and the randomized regions were subjected to a minimum Q-score threshold of 20. These reads were collected and written to a comma-separated values (csv) file. The graphs for number of barcodes/unique mutant and number of unique mutants/barcode were generated using bcpermut.py and aaRSperbc.py files, respectively. For more detailed instructions on using the NGS analysis pipeline, please refer to the Readme.md file in the link above.

**Total RNA isolation:**

DH10B cells were transformed with pBK lpp-tRNA$^{MaPyl}$-15AC1-metG-MaPylRS. Cells were grown overnight in 20 mL LB broth in the presence or absence of 1 mM ncAA and appropriate antibiotics. Cells were spun down at 5000 xg for 10 mins at 4 °C. The pellet was resuspended in 1 mL TRIzol™ reagent (Thermo Fisher) and incubated for 5 mins at RT. 200 µL of chloroform was added to the mixture, mixed thoroughly and incubated for 3 mins at RT. The solution was spun down at 12000 xg for 15 min at 4 °C. The clear supernatant was collected and mixed with 500 µL isopropanol. Then, the solution was incubated at 4 °C and spun down at 12000 xg for 10 min. The pellet was resuspended in 1 mL 70% ethanol and spun down at 7500 xg for 5 min. The pellet was air dried and resuspended in 50 µL dd$H_2O$. The total RNA concentration was determined using a Nanodrop spectrophotometer and the integrity was assessed with A260/A280.

**$NaIO_4$ oxidation:**

10 µg of total RNA was diluted up to 152 µL with oxidation buffer (50 mM NaOAc, 150 mM NaCl, 10 mM $MgCl_2$, 0.1 mM EDTA, pH 4.8) and 8 µL of 100 mM $NaIO_4$ (oxidation) or NaCl (no oxidation) was added[1]. The mixture was incubated at 25 °C for 60 mins and then quenched with 100 mM glucose for 10 mins at room temperature[2]. Finally, the product was recovered with ethanol precipitation by adding 400 µL of 100% ethanol and 1 µL glycogen (Thermo Scientific™ R0551). The mixture was spun down at 12,000

xg for 20 mins, followed by two washes with 95% ethanol. The pellet was air dried for 15 mins and resuspended in 40 µL $ddH_2O$.

**Deacylation**

tRNAs were deacylated with 100 mM Tris base (pH 9) and cleaned up with Zymo Research Oligo Clean and Concentrator kit (D4060) as per manufacturer's instructions. The product was eluted in 45 µL $ddH_2O$.

**tRNA-extension assay**

1 µL of 10 mM dNTPs, 1 µL of 1 µM tRNA-extension probe and 5 µL of NEBuffer 2 (B7002S) was added to the deacylated mixture. The mixture was subjected to a stepwise gradient: 95 °C, 70 °C, 50 °C for 2 min each and cooled to 4 °C. Subsequently, 0.5 µL of Klenow fragment (3' → 5' exo-) (M0212S) was added and the polymerase extension was carried out at 37 °C for 20 minutes[1]. 2x loading dye (4.8 g Urea, 30 µL Bromophenol blue, ~10 mL $ddH_2O$) was added to the mixture.

10% polyacrylamide 8M Urea denaturing gel was prepared using 14.4 g Urea, 10.3 mL 30% acrylamide/bisacrylamide solution, 37.5 :1 (Sigma-Aldrich A3699), and made up to 30 mL with 1x TBE. The solution was poured into gel cast after the addition of 200 µL 10% APS and 20 µL TEMED. The gels were pre-run at 250 V for 30 mins. 30 µL of the reaction mixture was run on the gel at 200 V for 90 mins and imaged on the Cy5 channel.

| **tREX probes** | **Sequence (5' to 3')** |
|---|---|
| 15A | AAAAAAAAAAAAAAA**TGGCGAGAGACCGGGGCGTCGAACCCCGC***|
| 19A2T5A3 | TTAAAAAAAAAAAAAAAAAAA**TGGCGAGAGACCGGGGCGTCGAACCCCGC**AAAAA* |
| 28A2T5A3 | TTAAAAAAAAAAAAAAAAAAAAAAAAAAAA**TGGCGAGAGACCGGGGCGTCGAACCCCGC**AAAAA* |

*indicates a Cy5 fluorophore on the 3'-end, red region indicates the extension sequence, **bold** indicates annealing to the tRNA

**Gel purification:**

The desired band was cut out using the Cy5 channel and transferred to a 1.5 mL microcentrifuge tube. To extract the tRNA, crush and soak method was used as described below. The bands were weighed and 1 µL of Crush and Soak buffer (CS buffer) was added per 2 mg. The gels were crushed into fine pieces with a squisher and 10 times more CS buffer was added (10 µL per 2 mg). The mixture was frozen at -80 °C for 15 min, thawed at 37°C for 15 min and incubated on a rocker at 4°C overnight. The samples were spun down at 12,000 xg for 15 mins at 4 °C and the supernatant containing tRNA (~400 µL) was ethanol precipitated (>2.5 h) as follows: 400 µL RNA, 800 µL 100% ethanol, 20 µL 3M NaOAc (pH 5.2), 1 µL glycogen. Finally, the mixture was spun down at 12,000 x g for 20 min at 4 °C, washed twice with 95% ethanol and eventually resuspended in 20 µL dd$H_2O$.

**Reverse transcription**

8 µL of the gel purified tRNA fraction was incubated with 1 µL of 10 mM dNTPs and 1 µL of 2.5 µM primer (Mapyl_polyT_R for adapter specific RT, MaPylT_RT_16nt_R for reverse transcription of $tRNA^{MaPyl}$) at 65 °C for 5 min. Further, 4 µL of 5x Induro® RT buffer and 1 µL of Induro® RT (NEB: M0681S) were added to the mixture and made up to 20 µL with dd$H_2O$. The reverse transcription was performed at 55 °C for 30 minutes and the enzyme was inactivated at 95 °C for 1 min. After cDNA synthesis, the tRNA was hydrolysed with 10 µL 1M NaOH, 20 µL 0.2 M EDTA (pH 8) at 65 °C for 15 min. The cDNA was recovered with Zymo Research Oligo Clean and Concentrator kit (D4060) and eluted in 20 µL dd$H_2O$.

**Amplification and sequencing**

The amplicon sequencing adapters were installed via PCR with primers QB_5_MaPylT_polyT_div_F and amplicon_tREX_15A_R. The NGS analysis for the selection outputs was performed using the following script: https://github.com/cpsoni74/Illumina_barcode_analysis.git[3]. Please refer to the NGS Analysis for Selection section for more details about the script.

**Illumina sequencing**

Illumina adapter sequences were attached to the DNA samples from the selection via two rounds of PCR. The primers were designed to add Illumina adapters provided by the TruSeq DNA HT Sample Prep Kit (Illumina). The forward adapter is AATGATACGGCGACCACCGAGATCTACAC[i5]ACACTCTTTCCCTACACGACGCTCTTCCGATCT, in which [i5] is an eight-nucleotide barcode sequence, and the reverse adapter is GATCGGAAGAGCACACGTCTGAACTCCAGTCAC[i7]ATCTCGTATGCCGTCTTCTGCTTG, in which [i7] is an eight-nucleotide barcode sequence. Both the i5 and i7 barcode sequences are provided below.

| Barcode | Sequence (5’ to 3’) |
|---|---|
| i5-D501 | TATAGCCT |
| i5-D502 | ATAGAGGC |
| i5-D503 | CCTATCCT |
| i5-D504 | GGCTCTGA |
| i7-D701 | CGAGTAAT |
| i7-D702 | TCTCCGGA |
| i7-D703 | AATGAGCG |
| i7-D704 | GGAATCTC |

The first set of primers, Ill-PylT-X1-F (X1a, X1b, X1c and X1d mixed in equimolar ratio) and Ill-PylT-X2-R (X2a, X2b, X2c and X2d mixed in equimolar ratio), consist of half of the TruSeq adapters, beginning immediately after the i5 or i7 barcode, followed by primer-binding sites to anneal to the sequences surrounding the tRNA library (for the forward primer: GGGGGACGGTCCGGCGAC; for the reverse primer: AAAAAAAAAAAAAAATGGCGAG). The second set of primers, a series of Illumina-i5-F and Illumina-i7-R variants containing different barcodes, consists of the 5′ half of the TruSeq adapters, followed by an i5 or i7 barcode, followed by primer-binding sites to anneal to the first PCR (for the forward primer: ACACTCTTTCCCTACACGACGC; for the reverse primer: GTGACTGGAGTTCAGACGTGTGCTC).

For the first PCR, samples were prepared using the primers Ill-PytlT-X1-F and Ill-PylT-X2-R and Phusion Hot Start II DNA Polymerase per the manufacturer’s instructions. PCR samples were purified with 1% agarose gel and a second round of PCR using Illumina-i5-F and Illumina-i7-R primer variants was performed to attach the region of the adapter sequences that includes the barcodes. Unique combinations of i5 and i7 barcode sequences were applied to each sample to enable multiplexing.

Samples were prepared for sequencing using the 150-cycle NextSeq500/550 Mid Output kit v2.5 (Illumina) according to the manufacturer’s instructions. Sequencing was carried out on an Illumina NextSeq500 System, with 40% Illumina PhiX Control.

**NGS Analysis for selection:**

The analysis was done using the following script: https://github.com/cpsoni74/Illumina_barcode_analysis.git. Please refer to the Readme.md file for detailed instructions on how the data was processed.

In summary, the reads were subjected to a Q-score filter (>30) and a mismatch filter, to discard any reads that are misaligned. Finally, the abundance and the fraction of total or relative abundance was calculated for each library member.

For each selection, three samples were sequenced. One for the input library (reference) and two for duplicates of the output library (selection). For each selection, the fold enrichment was determined by the ratio of relative abundance in output library vs input library. The enrichment factor was obtained by normalizing the fold enrichment of each member with respect to the most enriched member. Finally, each library member was ranked based on the average enrichment factor.

Only the tRNA barcodes present in the Pacbio sequencing data were collected and written to a csv file along with the associated MaPylRS mutants. The reads were further filtered such that the average enrichment in the presence of ncAA is at least 2-fold. Moreover, the difference in the average enrichment in +ncAA and -ncAA should be at least 2-fold. After filtering the reads, the tRNA barcodes were ranked based on the average enrichment in +ncAA selection. The average enrichment for each barcode in +ncAA and -ncAA selection were co-plotted on a scatter plot. The frequency plot weblogos were generated using https://weblogo.berkeley.edu/logo.cgi or using Logomaker.py

**Cloning hits from the selection:**

The MaPylRS hits from the selection were cloned using Gibson assembly. The hits were generated using overlap extension PCR similar to the MaPylRS library cloning using MaPylRS_NcoI_R_gib and MaPylRS_NdeI_F_gib. The final amplified insert was incubated with pBK-lpp-MaPylT-glnS-MaPylRS along with Gibson assembly HiFi mix at 55 °C for 60 min and transformed into DH10B electrocompetent cells. The hits were sequenced with Sanger Sequencing as well as whole plasmid sequencing.

**Assessment of MaPylRS mutants activity using sfGFP-TAG reporter:**

The pEvol-sfGFP-151-TAG reporter was co-transformed with a pBK plasmid containing $tRNA^{MaPyl}$ and MaPylRS variant into DH10B cells. Cultures were grown in 5 mL LB overnight and diluted to an OD 0.05 in 20 mL LB. Upon reaching an OD of 0.6, expression of the reporter was induced with 1 mM IPTG and appropriate ncAA. The cultures were incubated at 30 °C with shaking (250 rpm) for 16 hours. Subsequently, the cultures were spun down (4500 xg, 10 min, 4 °C), the media was removed and the pellet was resuspended in 1 mL 1x PBS. The resuspended cells were diluted 10-fold (15 µL in 135 µL PBS) and the fluorescence was measured in a black, clear-bottom 96-well plate using a plate reader (ex = 488 nm, em = 534 nm). Mean of two independent experiments were reported and the error bars represent standard deviation.

**sfGFP-151-TAG expression and purification:**

After inducing expression of a 20 mL culture similarly as described above, the cells were spun down (4500 xg, 10 min, 4 ℃), the media was removed and the pellets were resuspended in lysis buffer consisting of 1 mL B-Per™ (Thermo Scientific™ 78243), 10 µL Halt protease inhibitor cocktail (Thermo Scientific™ 78429) and 0.1 µL Pierce™ universal nuclease (Thermo Scientific™ 88702). The solution was incubated for 30 min at 4 ℃ and spun down at maximum speed. sfGFP was purified from the supernatant using HisPur™ Ni-NTA resin (Thermo Scientific™ 88221) as per manufacturer's instructions. Protein purity was characterized using SDS-PAGE and whole protein ESI-MS.

**Oligonucleotide sequences:**

| Oligo Name | Sequence (5' to 3') |
|---|---|
| lpp_BamHI_F | ATATAAAGGATCCCGCCGCTTCTTTGAGCGAACGATC |
| MaPylT_lib_R | ACCCGCTGGTCGCCGGACCGTCC |
| MaPyl_lib_10N_F | GGACGGTCCGGCGACCAGCGGGTNNNNNNNNNNACCTAGCCAGCGGGGTTCGACGC |
| tRNA_NotI_R | ATATATAGCGGCCGCGGATGAATGGCAGAAATTCGGTCGAC |
| MaPyl_15AC2_iR | CGTCGAACCCCGCTGGCTAGGTTTTCTAATCGTAGAGACCCGCTGGTCGCCGGACCG |
| MaPylT_15AC2_iF | ACCTAGCCAGCGGGGTTCGACG |
| MaPylT_15AC_F | GCGGGTCTCTAAACGATGGTAACCTAGCCAGCGGGGTTCGACGCCCCGGTCTCTCGCCACTGCAGATCCTTAGCGAAAG |

| | |
|---|---|
| MaPylT_15AC_R | CCCGCTGGCTAGGTTACCATCGTTTAGAGACCCGCTGGTCGCCGGACCGTCCCCCGAATTCAGCGTTACAAGTATTACACAAAG |
| Mapyl_polyT_R | AAAAAAAAAAAAAAATGGCGAG |
| MaPylT_RT_16nt_R | TGGCGAGAGACCGGGG |
| metG_NotI_F | ATATATAGCGGCCGCTTTACTTAACATTTTCCCATTTGGTACTATCTAACCCCTTTTCAC |
| MaPylRS_123_iR | GGAGCCAACATTGGGCGTAAGG |
| MaPylRS_L125NNK_iF | CGTGCCTTACGCCCAATGTTGGCTCCAAATNNKTATTCCGTTATGCGTGATTTGCGTGACCAC |
| MaPylRS_L165_iR | CAGCATCGTGAACTCCTCCAAATGCATGC |
| MaPylRS_N166, V168 NNK_iF | GGCATGCATTTGGAGGAGTTCACGATGCTGNNKCTTNNKGATATGGGACCGCGTGGTGATGCGACAGAGG |
| MaPylRS_NcoI_R | ATATATACCATGGTTAATTAATTTTTGCACCGTTCAGGTACGAGATACTTGCC |
| MaPylRS_125_V_R | CACGCAAATCACGCATAACGGAATACACATTTGGAGCCAACATTGGGCGTAAGG |
| MaPylRS_166_168_AV_R | GTCGCATCACCACGCGGTCCCATATCCACAAGCGCCAGCATCGTGAACTCCTCCAAATG |
| MaPylRS_166_168_AC_R | GTCGCATCACCACGCGGTCCCATATCGCAAAGCGCCAGCATCGTGAACTCCTCCAAATG |
| MaPylRS_166_168_SV_R | GTCGCATCACCACGCGGTCCCATATCCACAAGGCTCAGCATCGTGAACTCCTCCAAATG |
| pBK_MaPylRS_V_R_gib | GTGCGTCAGTGTATTTCACCGTCATATG |

| | |
|---|---|
| pBK_V_MaPylRS_gib_F | GGCAAGTATCTCGTACCTGAACGGTGCAAAAATTAATTAACCATGG |
| MaPylRS_NcoI_R_gib | CCATGGTTAATTAATTTTTGCACCGTTCAGGTACGAGATACTTGCC |
| MaPylRS_NdeI_F_gib | CATATGACGGTGAAATACACTGACGCAC |
| Ill_MaPylT_X1a_F | ACACTCTTTCCCTACACGACGCTCTTCCGATCTGGAACTCCTGGGGGACGGTCCGGCGAC |
| Ill_MaPylT_X1b_F | ACACTCTTTCCCTACACGACGCTCTTCCGATCTTAGCACATGGGGGGACGGTCCGGCGAC |
| Ill_MaPylT_X1c_F | ACACTCTTTCCCTACACGACGCTCTTCCGATCTCCTTGAGGAGGGGGACGGTCCGGCGAC |
| Ill_MaPylT_X1d_F | ACACTCTTTCCCTACACGACGCTCTTCCGATCTATCGTGTACGGGGGACGGTCCGGCGAC |
| Ill_MaPylT_X2a_R | TGGAGTTCAGACGTGTGCTCTTCCGATCTGACTACCCTAAAAAAAAAAAAAAATGGCGAG |
| Ill_MaPylT_X2b_R | TGGAGTTCAGACGTGTGCTCTTCCGATCTTCTGGATACAAAAAAAAAAAAAAATGGCGAG |
| Ill_MaPylT_X2c_R | TGGAGTTCAGACGTGTGCTCTTCCGATCTCTGATGGGAAAAAAAAAAAAAAAATGGCGAG |
| Ill_MaPylT_X2d_R | TGGAGTTCAGACGTGTGCTCTTCCGATCTAGACCTATGAAAAAAAAAAAAAAATGGCGAG |
| QB_5_MaPylT_polyT_div_F | TCCCTACACGACGCTCTTCCGATCTAAAATTTTTAAAAGGGGGACGGTCCGGCGACCAG |
| amplicon_tREX_15A_R | GACTGGAGTTCAGACGTGTGCTCTTCCGATCTAAAAAAAAAAAAAAATGGCGAG |

| | |
|---|---|
| Illumina-i5-D501-F | AATGATACGGCGACCACCGAGATCTACACTATAGCCTACACTCTTTCCCTACACGACGC |
| Illumina-i5-D502-F | AATGATACGGCGACCACCGAGATCTACACATAGAGGCACACTCTTTCCCTACACGACGC |
| Illumina-i5-D503-F | AATGATACGGCGACCACCGAGATCTACACCCTATCCTACACTCTTTCCCTACACGACGC |
| Illumina-i5-D504-F | AATGATACGGCGACCACCGAGATCTACACGGCTCTGAACACTCTTTCCCTACACGACGC |
| Illumina-i7-D701-R | CAAGCAGAAGACGGCATACGAGATCGAGTAATGTGACTGGAGTTCAGACGTGTGCTC |
| Illumina-i7-D702-R | CAAGCAGAAGACGGCATACGAGATTCTCCGGAGTGACTGGAGTTCAGACGTGTGCTC |
| Illumina-i7-D703-R | CAAGCAGAAGACGGCATACGAGATAATGAGCGGTGACTGGAGTTCAGACGTGTGCTC |
| Illumina-i7-D704-R | CAAGCAGAAGACGGCATACGAGATGGAATCTCGTGACTGGAGTTCAGACGTGTGCTC |

**Plasmid sequences:**

lpp promoter is highlighted in blue, tRNA$^{MaPyl}$ is highlighted in red, metG promoter is highlighted in purple, MaPylRS is highlighted in **brown**, glnS promoter is highlighted in orange, sfGFP is highlighted in green.

**pBK lpp-tRNA$^{MaPyl}$**

GGCTGGCCTGTTGAACAAGTCTGGAAAGAAATGCATAAGCTTTTGCCATTCTCACCGGATTC
AGTCGTCACTCATGGTGATTTCTCACTTGATAACCTTATTTTTGACGAGGGGAAATTAATAGG
TTGTATTGATGTTGGACGAGTCGGAATCGCAGACCGATACCAGGATCTTGCCATCCTATGGA
ACTGCCTCGGTGAGTTTTCTCCTTCATTACAGAAACGGCTTTTTCAAAAATATGGTATTGATA
ATCCTGATATGAATAAATTGCAGTTTCATTTGATGCTCGATGAGTTTTTCTAATCAGAATTGG
TTAATTGGTTGTAACACTGGCAGAGCATTACGCTGACTTGACGGGACGGCGGCTTTGTTGAA
TAAATCGAACTTTTGCTGAGTTGAAGGATCCCGCCGCTTCTTTGAGCGAACGATCAAAAATA
AGTGGCGCCCCATCAAAAAAATATTCTCAACATAAAAAACTTTGTGTAATACTTGTAACGCT
GAATTCGGGGGACGGTCCGGCGACCAGCGGGTCTCTAAAACCTAGCCAGCGGGGTTCGACG
CCCCGGTCTCTCGCCACTGCAGATCCTTAGCGAAAGCTAAGGATTTTTTTAGTCGACCGAAT
TTCTGCCATTCATCCGCGGCCGCCTGCAGTTTCAAACGCTAAATTGCCTGATGCGCTACGCTT
ATCAGGCCTACATGATCTCTGCAATATATTGAGTTTGCGTGCTTTTGTAGGCCGGATAAGGC
GTTCACGCCGCATCCGGCAAGAAACAGCAAACAATCACATGTGAGCAAAAGGCCAGCAAAA
GGCCAGGAACCGTAAAAAGGCCGCGTTGCTGGCGTTTTTCCATAGGCTCCGCCCCCCTGACG
AGCATCACAAAAATCGACGCTCAAGTCAGAGGTGGCGAAACCCGACAGGACTATAAAGATA
CCAGGCGTTTCCCCCTGGAAGCTCCCTCGTGCGCTCTCCTGTTCCGACCCTGCCGCTTACCGG
ATACCTGTCCGCCTTTCTCCCTTCGGGAAGCGTGGCGCTTTCTCATAGCTCACGCTGTAGGTA
TCTCAGTTCGGTGTAGGTCGTTCGCTCCAAGCTGGGCTGTGTGCACGAACCCCCCGTTCAGC
CCGACCGCTGCGCCTTATCCGGTAACTATCGTCTTGAGTCCAACCCGGTAAGACACGACTTA
TCGCCACTGGCAGCAGCCACTGGTAACAGGATTAGCAGAGCGAGGTATGTAGGCGGTGCTA
CAGAGTTCTTGAAGTGGTGGCCTAACTACGGCTACACTAGAAGGACAGTATTTGGTATCTGC
GCTCTGCTGAAGCCAGTTACCTTCGGAAAAAGAGTTGGTAGTTCTTGATCCGGCAAACAAAC
CACCGCTGGTAGCGGTGGTTTTTTTGTTTGCAAGCAGCAGATTACGCGCAGAAAAAAAGGAT
CTCAAGAAGATCCTTTGATCTTTTCTACGGGGTCTGACGCTCAGTGGAACGAAAACTCACGT
TAAGGGATTTTGGTCATGAACAATAAAACTGTCTGCTTACATAAACAGTAATACAAGGGGTG
TTATGAGCCATATTCAACGGGAAACGTCTTGCTCGAGGCCGCGATTAAATTCCAACATGGAT
GCTGATTTATATGGGTATAAATGGGCTCGCGATAATGTCGGGCAATCAGGTGCGACAATCTA
TCGATTGTATGGGAAGCCCGATGCGCCAGAGTTGTTTCTGAAACATGGCAAAGGTAGCGTTG
CCAATGATGTTACAGATGAGATGGTCAGACTAAACTGGCTGACGGAATTTATGCCTCTTCCG
ACCATCAAGCATTTTATCCGTACTCCTGATGATGCATGGTTACTCACCACTGCGATCCCCGGG
AAAACAGCATTCCAGGTATTAGAAGAATATCCTGATTCAGGTGAAAATATTGTTGATGCGCT
GGCAGTGTTCCTGCGCCGGTTGCATTCGATTCCTGTTTGTAATTGTCCTTTTAACAGCGATCG
CGTATTTCGTCTCGCTCAGGCGCAATCACGAATGAATAACGGTTTGGTTGATGCGAGTGATT
TTGATGACGAGCGTAAT

**pBK lpp -tRNA$^{MaPyl}$-metG-MaPylRS**

GTCGACCGAATTTCTGCCATTCATCCGCGGCCGCTT**TACTTAACATTTTCCCATTTGGTACT**
**ATCTAACCCCTTTTCACTATTAAGAAGTAATGCCTACT**CAT**ATGACGGTGAAATACACT**
**GACGCACAGATCCAGCGCCTTCGCGAATATGGGAATGGCACGTATGAACAGAAAGTGT**
**TCGAAGATTTGGCTTCGCGCGACGCAGCCTTTAGCAAAGAAATGAGTGTTGCCTCAAC**
**CGACAATGAGAAAAAAATTAAGGGCATGATTGCCAACCCGTCACGTCATGGACTTACG**
**CAACTTATGAACGACATTGCCGACGCATTAGTCGCTGAGGGATTTATCGAGGTCCGCA**
**CGCCAATCTTTATCTCAAAAGACGCGCTTGCCCGTATGACGATTACAGAAGACAAGCC**
**CCTGTTCAAGCAAGTATTCTGGATCGACGAGAAGCGTGCCTTACGCCCAATGTTGGCT**
**CCAAATTTATATTCCGTTATGCGTGATTTGCGTGACCACACCGACGGCCCAGTGAAGAT**
**TTTCGAGATGGGGAGCTGTTTTCGCAAGGAAAGTCACAGTGGCATGCATTTGGAGGAG**
**TTCACGATGCTGAACCTTGTGGATATGGGACCGCGTGGTGATGCGACAGAGGTTTTAA**
**AAAATTACATTAGTGTTGTGATGAAAGCAGCGGGATTGCCCGATTATGATTTAGTCCAG**
**GAAGAGAGTGACGTCTACAAAGAAACTATCGATGTTGAGATTAACGGGCAAGAAGTAT**
**GTAGCGCTGCTGTCGGACCCCATTATCTGGATGCTGCCCATGATGTGCATGAACCTTG**
**GTCTGGTGCTGGTTTCGGTTTGGAGCGCTTATTAACCATTCGTGAGAAATATTCCACAG**
**TAAAGAAAGGGGGGGCAAGTATCTCGTACCTGAACGGTGCAAAAATTAATTAA**CCATG
GCTGCAGTTTCAAACGCTAAATTGCCTGATGCGCTACGCTTATCAGGCCTACATGATCTCTGC
AATATATTGAGTTTGCGTGCTTTTGTAGGCCGGATAAGGCGTTCACGCCGCATCCGGCAAGA
AACAGCAAACAATCCAAAACGCCGCGTTCAGCGGCGTTTTTTCTGCTTTTCTTCGCGAATTA
ATTCCGCTTCGCACATGTGAGCAAAAGGCCAGCAAAAGGCCAGGAACCGTAAAAAGGCCGC
GTTGCTGGCGTTTTTCCATAGGCTCCGCCCCCCTGACGAGCATCACAAAAATCGACGCTCAA
GTCAGAGGTGGCGAAACCCGACAGGACTATAAAGATACCAGGCGTTTCCCCCTGGAAGCTC
CCTCGTGCGCTCTCCTGTTCCGACCCTGCCGCTTACCGGATACCTGTCCGCCTTTCTCCCTTCG
GGAAGCGTGGCGCTTTCTCATAGCTCACGCTGTAGGTATCTCAGTTCGGTGTAGGTCGTTCG
CTCCAAGCTGGGCTGTGTGCACGAACCCCCCGTTCAGCCCGACCGCTGCGCCTTATCCGGTA
ACTATCGTCTTGAGTCCAACCCGGTAAGACACGACTTATCGCCACTGGCAGCAGCCACTGGT
AACAGGATTAGCAGAGCGAGGTATGTAGGCGGTGCTACAGAGTTCTTGAAGTGGTGGCCTA
ACTACGGCTACACTAGAAGGACAGTATTTGGTATCTGCGCTCTGCTGAAGCCAGTTACCTTC
GGAAAAAGAGTTGGTAGTTCTTGATCCGGCAAACAAACCACCGCTGGTAGCGGTGGTTTTTT
TGTTTGCAAGCAGCAGATTACGCGCAGAAAAAAAGGATCTCAAGAAGATCCTTTGATCTTTT
CTACGGGGTCTGACGCTCAGTGGAACGAAAACTCACGTTAAGGGATTTTGGTCATGAACAAT
AAAACTGTCTGCTTACATAAACAGTAATACAAGGGGTGTTATGAGCCATATTCAACGGGAA
ACGTCTTGCTCGAGGCCGCGATTAAATTCCAACATGGATGCTGATTTATATGGGTATAAATG
GGCTCGCGATAATGTCGGGCAATCAGGTGCGACAATCTATCGATTGTATGGGAAGCCCGATG
CGCCAGAGTTGTTTCTGAAACATGGCAAAGGTAGCGTTGCCAATGATGTTACAGATGAGATG
GTCAGACTAAACTGGCTGACGGAATTTATGCCTCTTCCGACCATCAAGCATTTTATCCGTACT
CCTGATGATGCATGGTTACTCACCACTGCGATCCCCGGAAAACAGCATTCCAGGTATTAGA
AGAATATCCTGATTCAGGTGAAAATATTGTTGATGCGCTGGCAGTGTTCCTGCGCCGGTTGC
ATTCGATTCCTGTTTGTAATTGTCCTTTTAACAGCGATCGCGTATTTCGTCTCGCTCAGGCGC
AATCACGAATGAATAACGGTTTGGTTGATGCGAGTGATTTTGATGACGAGCGTAATGGCTGG
CCTGTTGAACAAGTCTGGAAAGAAATGCATAAGCTTTTGCCATTCTCACCGGATTCAGTCGT
CACTCATGGTGATTTCTCACTTGATAACCTTATTTTTGACGAGGGGAAATTAATAGGTTGTAT
TGATGTTGGACGAGTCGGAATCGCAGACCGATACCAGGATCTTGCCATCCTATGGAACTGCC
TCGGTGAGTTTTCTCCTTCATTACAGAAACGGCTTTTTCAAAAATATGGTATTGATAATCCTG

ATATGAATAAATTGCAGTTTCATTTGATGCTCGATGAGTTTTTCTAATCAGAATTGGTTAATT
GGTTGTAACACTGGCAGAGCATTACGCTGACTTGACGGGACGGCGGCTTTGTTGAATAAATC
GAACTTTTGCTGAGTTGAAGGATCCCGCCGCTTCTTTGAGCGAACGATCAAAAATAAGTGGC
GCCCCATCAAAAAAATATTCTCAACATAAAAAACTTTGTGTAATACTTGTAACGCTGAATTC
GGGGGACGGTCCGGCGACCAGCGGGTCTCTAAAACCTAGCCAGCGGGGTTCGACGCCCCGG
TCTCTCGCCACTGCAGATCCTTAGCGAAAGCTAAGGATTTTTTTA

**pBK lpp-tRNA$^{MaPyl}$-glnS-MaPylRS**

GTCGACCGAATTTCTGCCATTCATCCGCGGCCGCTCGGGTTGTCAGCCTGTCCCGCTTATAAG
ATCATACGCCGTTATACGTTGTTTACGCTTTGAGGAATCCCAT**ATGACGGTGAAATACACT**
**GACGCACAGATCCAGCGCCTTCGCGAATATGGGAATGGCACGTATGAACAGAAAGTGT**
**TCGAAGATTTGGCTTCGCGCGACGCAGCCTTTAGCAAAGAAATGAGTGTTGCCTCAAC**
**CGACAATGAGAAAAAAATTAAGGGCATGATTGCCAACCCGTCACGTCATGGACTTACG**
**CAACTTATGAACGACATTGCCGACGCATTAGTCGCTGAGGGATTTATCGAGGTCCGCA**
**CGCCAATCTTTATCTCAAAAGACGCGCTTGCCCGTATGACGATTACAGAAGACAAGCC**
**CCTGTTCAAGCAAGTATTCTGGATCGACGAGAAGCGTGCCTTACGCCCAATGTTGGCT**
**CCAAATTTATATTCCGTTATGCGTGATTTGCGTGACCACACCGACGGCCCAGTGAAGAT**
**TTTCGAGATGGGGAGCTGTTTTCGCAAGGAAAGTCACAGTGGCATGCATTTGGAGGAG**
**TTCACGATGCTGAACCTTGTGGATATGGGACCGCGTGGTGATGCGACAGAGGTTTTAA**
**AAAATTACATTAGTGTTGTGATGAAAGCAGCGGGATTGCCCGATTATGATTTAGTCCAG**
**GAAGAGAGTGACGTCTACAAAGAAACTATCGATGTTGAGATTAACGGGCAAGAAGTAT**
**GTAGCGCTGCTGTCGGACCCCATTATCTGGATGCTGCCCATGATGTGCATGAACCTTG**
**GTCTGGTGCTGGTTTCGGTTTGGAGCGCTTATTAACCATTCGTGAGAAATATTCCACAG**
**TAAAGAAAGGGGGGGCAAGTATCTCGTACCTGAACGGTGCAAAAATTAATTAA**CCATG
GCTGCAGTTTCAAACGCTAAATTGCCTGATGCGCTACGCTTATCAGGCCTACATGATCTCTGC
AATATATTGAGTTTGCGTGCTTTTGTAGGCCGGATAAGGCGTTCACGCCGCATCCGGCAAGA
AACAGCAAACAATCCAAAACGCCGCGTTCAGCGGCGTTTTTTCTGCTTTTCTTCGCGAATTA
ATTCCGCTTCGCACATGTGAGCAAAAGGCCAGCAAAAGGCCAGGAACCGTAAAAAGGCCGC
GTTGCTGGCGTTTTTCCATAGGCTCCGCCCCCCTGACGAGCATCACAAAAATCGACGCTCAA
GTCAGAGGTGGCGAAACCCGACAGGACTATAAAGATACCAGGCGTTTCCCCCTGGAAGCTC
CCTCGTGCGCTCTCCTGTTCCGACCCTGCCGCTTACCGGATACCTGTCCGCCTTTCTCCCTTCG
GGAAGCGTGGCGCTTTCTCATAGCTCACGCTGTAGGTATCTCAGTTCGGTGTAGGTCGTTCG
CTCCAAGCTGGGCTGTGTGCACGAACCCCCCGTTCAGCCCGACCGCTGCGCCTTATCCGGTA
ACTATCGTCTTGAGTCCAACCCGGTAAGACACGACTTATCGCCACTGGCAGCAGCCACTGGT
AACAGGATTAGCAGAGCGAGGTATGTAGGCGGTGCTACAGAGTTCTTGAAGTGGTGGCCTA
ACTACGGCTACACTAGAAGGACAGTATTTGGTATCTGCGCTCTGCTGAAGCCAGTTACCTTC
GGAAAAAGAGTTGGTAGTTCTTGATCCGGCAAACAAACCACCGCTGGTAGCGGTGGTTTTTT
TGTTTGCAAGCAGCAGATTACGCGCAGAAAAAAAGGATCTCAAGAAGATCCTTTGATCTTTT
CTACGGGGTCTGACGCTCAGTGGAACGAAAACTCACGTTAAGGGATTTTGGTCATGAACAAT
AAAACTGTCTGCTTACATAAACAGTAATACAAGGGGTGTTATGAGCCATATTCAACGGGAA
ACGTCTTGCTCGAGGCCGCGATTAAATTCCAACATGGATGCTGATTTATATGGGTATAAATG
GGCTCGCGATAATGTCGGGCAATCAGGTGCGACAATCTATCGATTGTATGGGAAGCCCGATG
CGCCAGAGTTGTTTCTGAAACATGGCAAAGGTAGCGTTGCCAATGATGTTACAGATGAGATG
GTCAGACTAAACTGGCTGACGGAATTTATGCCTCTTCCGACCATCAAGCATTTTATCCGTACT

CCTGATGATGCATGGTTACTCACCACTGCGATCCCCGGGAAAACAGCATTCCAGGTATTAGA
AGAATATCCTGATTCAGGTGAAAATATTGTTGATGCGCTGGCAGTGTTCCTGCGCCGGTTGC
ATTCGATTCCTGTTTGTAATTGTCCTTTTAACAGCGATCGCGTATTTCGTCTCGCTCAGGCGC
AATCACGAATGAATAACGGTTTGGTTGATGCGAGTGATTTTGATGACGAGCGTAATGGCTGG
CCTGTTGAACAAGTCTGGAAAGAAATGCATAAGCTTTTGCCATTCTCACCGGATTCAGTCGT
CACTCATGGTGATTTCTCACTTGATAACCTTATTTTTGACGAGGGGAAATTAATAGGTTGTAT
TGATGTTGGACGAGTCGGAATCGCAGACCGATACCAGGATCTTGCCATCCTATGGAACTGCC
TCGGTGAGTTTTCTCCTTCATTACAGAAACGGCTTTTTCAAAAATATGGTATTGATAATCCTG
ATATGAATAAATTGCAGTTTCATTTGATGCTCGATGAGTTTTTCTAATCAGAATTGGTTAATT
GGTTGTAACACTGGCAGAGCATTACGCTGACTTGACGGGACGGCGGCTTTGTTGAATAAATC
GAACTTTTGCTGAGTTGAAGGATCCCGCCGCTTCTTTGAGCGAACGATCAAAAATAAGTGGC
GCCCCATCAAAAAAATATTCTCAACATAAAAAACTTTGTGTAATACTTGTAACGCTGAATTC
GGGGGACGGTCCGGCGACCAGCGGGTCTCTAAAACCTAGCCAGCGGGGTTCGACGCCCCGG
TCTCTCGCCACTGCAGATCCTTAGCGAAAGCTAAGGATTTTTTTA

pEvol T5-lac-sfGFP-151-TAG:

CTCCATTTTAGCTTCCTTAGCTCCTGAAAATCTCGATAACTCAAAAAATACGCCCGGTAGTG
ATCTTATTTCATTATGGTGAAAGTTGGAACCTCTTACGTGCCGATCAACGTCTCATTTTCGCC
AAAAGTTGGCCCAGGGCTTCCCGGTATCAACAGGGACACCAGGATTTATTTATTCTGCGAAG
TGATCTTCCGTCACAGGTATTTGTTCGGCGCAAAGTGCGTCGGGTGATGCTGCCAACTTACT
GATTTAGTGTATGATGGTGTTTTTGAGGTGCTCCAGTGGCTTCTGTTTCTATCAGCTGTCCCT
CCTGTTCAGCTACTGACGGGGTGGTGCGTAACGGCAAAAGCACCGCCGGACATCAGCGCTA
GCGGAGTGTATACTGGCTTACTATGTTGGCACTGATGAGGGTGTCAGTGAAGTGCTTCATGT
GGCAGGAGAAAAAAGGCTGCACCGGTGCGTCAGCAGAATATGTGATACAGGATATATTCCG
CTTCCTCGCTCACTGACTCGCTACGCTCGGTCGTTCGACTGCGGCGAGCGGAAATGGCTTAC
GAACGGGGCGGAGATTTCCTGGAAGATGCCAGGAAGATACTTAACAGGGAAGTGAGAGGG
CCGCGGCAAAGCCGTTTTTCCATAGGCTCCGCCCCCCTGACAAGCATCACGAAATCTGACGC
TCAAATCAGTGGTGGCGAAACCCGACAGGACTATAAAGATACCAGGCGTTTCCCCCTGGCG
GCTCCCTCGTGCGCTCTCCTGTTCCTGCCTTTCGGTTTACCGGTGTCATTCCGCTGTTATGGCC
GCGTTTGTCTCATTCCACGCCTGACACTCAGTTCCGGGTAGGCAGTTCGCTCCAAGCTGGACT
GTATGCACGAACCCCCCGTTCAGTCCGACCGCTGCGCCTTATCCGGTAACTATCGTCTTGAGT
CCAACCCGGAAAGACATGCAAAAGCACCACTGGCAGCAGCCACTGGTAATTGATTTAGAGG
AGTTAGTCTTGAAGTCATGCGCCGGTTAAGGCTAAACTGAAAGGACAAGTTTTGGTGACTGC
GCTCCTCCAAGCCAGTTACCTCGGTTCAAAGAGTTGGTAGCTCAGAGAACCTTCGAAAAACC
GCCCTGCAAGGCGGTTTTTTCGTTTTCAGAGCAAGAGATTACGCGCAGACCAAAACGATCTC
AAGAAGATCATCTTATTAATCAGATAAAATATTTCTAGATTTCAGTGCAATTTATCTCTTCAA
ATGTAGCACCTGAAGTCAGCCCCATACGATATAAGTTGTAATTCTCATGTTTGACAGCTTATC
ATCGATAAGCTTGCAATTTATCTCTTCAAATGTAGCACCTGAAGTCAGCCCCATACGATATA
AGTTGTAATTCTCATGTTAGTCATGCCCCGCGCCCACCGGAAGGAGCTGACTGGGTTGAAGG
CTCTCAAGGGCATCGGTCGAGATCCCGGTGCCTAATGAGTGAGCTAACTTACATTAATTGCG
TTGCGCTCACTGCCCGCTTTCCAGTCGGGAAACCTGTCGTGCCAGCTGCATTAATGAATCGG
CCAACGCGCGGGGAGAGGCGGTTTGCGTATTGGGCGCCAGGGTGGTTTTTCTTTTCACCAGT
GAGACGGGCAACAGCTGATTGCCCTTCACCGCCTGGCCCTGAGAGAGTTGCAGCAAGCGGT
CCACGCTGGTTTGCCCCAGCAGGCGAAAATCCTGTTTGATGGTGGTTAACGGCGGGATATAA

CATGAGCTGTCTTCGGTATCGTCGTATCCCACTACCGAGATGTCCGCACCAACGCGCAGCCC
GGACTCGGTAATGGCGCGCATTGCGCCCAGCGCCATCTGATCGTTGGCAACCAGCATCGCAG
TGGGAACGATGCCCTCATTCAGCATTTGCATGGTTTGTTGAAAACCGGACATGGCACTCCAG
TCGCCTTCCCGTTCCGCTATCGGCTGAATTTGATTGCGAGTGAGATATTTATGCCAGCCAGCC
AGACGCAGACGCGCCGAGACAGAACTTAATGGGCCCGCTAACAGCGCGATTTGCTGGTGAC
CCAATGCGACCAGATGCTCCACGCCCAGTCGCGTACCGTCTTCATGGGAGAAAATAATACTG
TTGATGGGTGTCTGGTCAGAGACATCAAGAAATAACGCCGGAACATTAGTGCAGGCAGCTT
CCACAGCAATGGCATCCTGGTCATCCAGCGGATAGTTAATGATCAGCCCACTGACGCGTTGC
GCGAGAAGATTGTGCACCGCCGCTTTACAGGCTTCGACGCCGCTTCGTTCTACCATCGACAC
CACCACGCTGGCACCCAGTTGATCGGCGCGAGATTTAATCGCCGCGACAATTTGCGACGGCG
CGTGCAGGGCCAGACTGGAGGTGGCAACGCCAATCAGCAACGACTGTTTGCCCGCCAGTTG
TTGTGCCACGCGGTTGGGAATGTAATTCAGCTCCGCCATCGCCGCTTCCACTTTTTCCCGCGT
TTTCGCAGAAACGTGGCTGGCCTGGTTCACCACGCGGGAAACGGTCTGATAAGAGACACCG
GCATACTCTGCGACATCGTATAACGTTACTGGTTTCACATTCACCACCCTGAATTGACTCTCT
TCCGGGCGCTATCATGCCATACCGCGAAAGGTTTTGCGCCATTCGATGGTGTCCGGGATCTC
GACGCTCTCCCTTATGCGACTCCTGCATTAGGCTCACTATAGGGGAATTGTGAGCGGATAAC
AATTCCCCTCTAGAGTTTGACAGCATTGTCATCGATCTCGAGAAATCATAAAAAATTTATTTG
CTTTGTGAGCGGATAACAATTATAATAGATTCAATTGTGAGCGGATAACAATTTCACACAGA
ATTCATTAAAGAGGAGAAATTACATATGAGCAAAGGAGAAGAACTTTTCACTGGAGTTGTC
CCAATTCTTGTTGAATTAGATGGTGATGTTAATGGGCACAAATTTTCTGTCCGTGGAGAGGG
TGAAGGTGATGCTACAAACGGAAAACTCACCCTTAAATTTATTTGCACTACTGGAAAACTAC
CTGTTCCGTGGCCAACACTTGTCACTACTCTGACCTATGGTGTTCAATGCTTTTCCCGTTATC
CGGATCACATGAAACGGCATGACTTTTTCAAGAGTGCCATGCCCGAAGGTTATGTACAGGAA
CGCACTATATCTTTCAAAGATGACGGGACCTACAAGACGCGTGCTGAAGTCAAGTTTGAAGG
TGATACCCTTGTTAATCGTATCGAGTTAAAGGGTATTGATTTTAAAGAAGATGGAAACATTC
TTGGACACAAACTCGAGTACAACTTTAACTCACACAATGTATAGATCACGGCAGACAAACA
AAAGAATGGAATCAAAGCTAACTTCAAAATTCGCCACAACGTTGAAGATGGTTCCGTTCAAC
TAGCAGACCATTATCAACAAAATACTCCAATTGGCGATGGCCCTGTCCTTTTACCAGACAAC
CATTACCTGTCGACACAATCTGTCCTTTCGAAAGATCCCAACGAAAAGCGTGACCACATGGT
CCTTCTTGAGTTTGTAACTGCTGCTGGGATTACACATGGCATGGATGAGCTCTACAAAGGAT
CCCACCACCACCACCACCACTAAAAGCTTAATTAGCTGAGCTTGGACTCCTGTTGATAGATC
CAGTAATGACCTCAGAACTCCATCTGGATTTGTTCAGAACGCTCGGTTGCCGCCGGGCGTTT
TTTATTGGTGAGAATCCAAGCTAGCTTGGCGCCTCGAGCAGCTCAGGGTCGAATTTGCTTTC
GAATTTCTGCCATTCATCCGCTTATTATCACTTATTCAGGCGTAGCAACCAGGCGTTTAAGGG
CACCAATAACTGCCTTAAAAAAATTACGCCCCGCCCTGCCACTCATCGCAGTACTGTTGTAA
TTCATTAAGCATTCTGCCGACATGGAAGCCATCACAAACGGCATGATGAACCTGAATCGCCA
GCGGCATCAGCACCTTGTCGCCTTGCGTATAATATTTGCCCATGGTGAAAACGGGGGCGAAG
AAGTTGTCCATATTGGCCACGTTTAAATCAAAACTGGTGAAACTCACCCAGGGATTGGCTGA
GACGAAAAACATATTCTCAATAAACCCTTTAGGGAAATAGGCCAGGTTTTCACCGTAACACG
CCACATCTTGCGAATATATGTGTAGAAACTGCCGGAAATCGTCGTGGTATTCACTCCAGAGC
GATGAAAACGTTTCAGTTTGCTCATGGAAAACGGTGTAACAAGGGTGAACACTATCCCATAT
CACCAGCTCACCTTCTTTCATTGCCATACGGAATTCCGGATGAGCATTCATCAGGCGGGCAA
GAATGTGAATAAAGGCCGGATAAAACTTGTGCTTATTTTTCTTTACGGTCTTTAAAAAGGCC
GTAATATCCAGCTGAACGGTCTGGTTATAGGTACATTGAGCAACTGACTGAAATGCCTCAAA
ATGTTCTTTACGATGCCATTGGGATATATCAACGGTGGTATATCCAGTGATTTTTTT

**References**:


1. Cervettini, D., Tang, S., Fried, S. D., Willis, J. C. W., Funke, L. F. H., Colwell, L. J., and Chin, J. W. (2020) Rapid discovery and evolution of orthogonal aminoacyl-tRNA synthetase–tRNA pairs, *Nature Biotechnology 38*, 989-999.
2. Passarelli, M. C., Pinzaru, A. M., Asgharian, H., Liberti, M. V., Heissel, S., Molina, H., Goodarzi, H., and Tavazoie, S. F. (2022) Leucyl-tRNA synthetase is a tumour suppressor in breast cancer and regulates codon-dependent translation dynamics, *Nature Cell Biology 24*, 307-315.
3. Jewel, D., Kelemen, R. E., Huang, R. L., Zhu, Z., Sundaresh, B., Cao, X., Malley, K., Huang, Z., Pasha, M., Anthony, J., van Opijnen, T., and Chatterjee, A. (2023) Virus-assisted directed evolution of enhanced suppressor tRNAs in mammalian cells, *Nature methods 20*, 95-103.
